# A novel rhesus macaque model of Huntington’s disease recapitulates key neuropathological changes along with progressive motor and cognitive decline

**DOI:** 10.1101/2022.02.02.478920

**Authors:** Alison R. Weiss, William A. Liguore, Kristin Brandon, Xiaojie Wang, Zheng Liu, Jacqueline S. Domire, Dana Button, Christopher D. Kroenke, Jodi L. McBride

## Abstract

We created a new nonhuman primate model of the genetic neurodegenerative disorder, Huntington’s disease (HD), by injecting a mixture of recombinant adeno-associated viral vectors, serotypes AAV2 and AAV2.retro, each expressing a fragment of human mutant *HTT* (*mHTT)* into the caudate and putamen of adult rhesus macaques. This novel modeling strategy results in robust expression of mutant huntingtin protein (mHTT) in the injected brain regions, as well as dozens of other cortical and subcortical brain regions that are also affected in human HD patients. We queried the disruption of cortico-basal ganglia circuitry for 20-months post-surgery using a variety of behavioral and imaging readouts. Compared to controls, mHTT-treated macaques developed progressive working memory decline and motor impairment. Multimodal imaging revealed circuit-wide white and gray matter degenerative processes in several key brain regions affected in HD. This novel model will aid in the development of disease biomarkers and therapeutic strategies for this devastating disorder.

## INTRODUCTION

Huntington’s disease (HD) is a genetic, progressive neurodegenerative disorder caused by an expanded CAG repeat in exon 1 of the *HTT* gene ^1^. When the glutamine stretch in the N-terminus of huntingtin protein (HTT) exceeds approximately 40 repeats, HTT misfolds and sets off a toxic sequence of events inside the cell including transcriptional dysregulation, mitochondrial dysfunction, calcium signaling disruption and altered neurotransmission, including glutamatergic and dopaminergic dysregulation ^2^. Accordingly, post-mortem studies of HD patient brain tissue have shown significant neuropathology including mutant huntingtin protein (mHTT) aggregate formation, profound neuronal death and gliosis ^3–5^. The most highly affected regions are the caudate nucleus and putamen (together comprising the striatum); however numerous brain regions that send afferent connections to the striatum are also affected, including widespread cortical and subcortical brain areas ^6–10^. The network of white matter fiber tracts interconnecting the striatum with the cortex also show extensive evidence of degeneration ^11–14^, highlighting that HD is not only a striatal disorder, but rather a disease characterized by a progressive disintegration of cortico-basal ganglia circuitry.

The symptoms of HD are extremely detrimental to quality of life, as patients suffer from a progressive movement disorder that is characterized by lack of balance and coordination, altered fine motor skills and hyperkinetic involuntary movements of the limbs, trunk and face, known as chorea. In later stages of the disease patients can experience bradykinetic movements, dystonia and rigidity ^15–17^. These motor phenotypes are often accompanied by deteriorating cognitive function, including working memory decline and a reduced capacity to plan and organize daily tasks. HD patients also exhibit profound personality changes and mood disturbances that can include depression, anxiety and irritability ^18–22^. Currently there are no treatments capable of slowing disease progression. Therefore, to aid in the creation and evaluation of effective therapeutic interventions, it is critical to develop appropriate animal models of HD.

Genetically modified rodents have been key in advancing our understanding of pathophysiological mechanisms that are altered by the *mHTT* gene, and numerous mouse and rat models have been created over the past three decades, reviewed in Farshim and Bates, 2018 ^23^. These models have played a critical role in characterizing HTT-mediated disease pathology, and several have been utilized to screen therapeutic strategies. However, an important consideration is that these models do not fully recapitulate the behavioral symptomology of humans with HD. Some HD mouse models display balance issues that can be measured via rotorod performance ^24^, but do not display the hallmark signs of chorea, bradykinesia nor fine motor skill deficits. Similarly, some mouse models exhibit spatial memory deficits but lack other signature features of HD^24^. Furthermore, species differences in genetics, brain size, structure and neural connectivity all limit the ability to translate findings from rodents to predict clinical responses in human patients. For example, there still exists an ambiguity on whether specific frontal cortical areas in monkeys and humans, such as the medial and lateral prefrontal cortices (areas known to be affected in HD) share cross-species homologies with rodents ^25^. Perhaps most striking to consider is that none of the therapies that have shown efficacy in mouse models have resulted in success in clinical trials over the past several decades. For these reasons, large animal models including sheep, minipigs and nonhuman primates (NHP) have emerged more recently as a new and critical tool for the HD research community ^26^.

The earliest NHP models of HD were created by using neurotoxins to lesion the caudate and/or putamen in order to replicate the profound striatal atrophy and hyperkinetic movements seen in patients ^26^. However, the toxin-based approach decreased in popularity after the *HTT* gene mutation was identified. Subsequent NHP models have been created by viral-mediated delivery of a fragment of the human mHTT gene ^27, 28^ or via the development of transgenic HD macaques ^29^, both bearing *HTT* genes with expanded CAG repeats that encode mHTT proteins with elongated polyglutamine tracts (Q) at the N terminus. Palfi and colleagues created the first viral-mediated NHP model of HD by injecting a lentiviral vector expressing a fragment of *HTT* with 82 CAG repeats (LV-HTT82Q) bilaterally into the dorsolateral, posterior portion of putamen of three rhesus macaques ^27^. LV-HTT82Q macaques developed dyskinetic movements of the limbs and trunk beginning at 15 weeks post-surgery and persisting out to 30-weeks post-surgery. Brain tissue revealed mHTT inclusion formation, neuronal loss and astrocytosis in the circumscribed area of injection. Given that HD pathology in patients has been identified throughout the putamen, caudate nucleus, and in multiple other cortical and sub-cortical brain regions, it is essential to create a virally-mediated NHP model that more closely recapitulates this widespread pattern of neurodegeneration and circuit dysfunction. Moreover, because cognitive dysfunction dramatically alters the HD patient’s quality of life and autonomy, an NHP model that exhibits both motor and working memory decline will be more useful for evaluating promising therapeutics.

Recent techniques using directed evolution approaches to develop new adeno-associated viruses (AAV) have produced novel capsid variants that exhibit enhanced properties of retrograde and/or anterograde transport in brains of both rodents and NHPs ^28, 30–32^. Utilizing these new AAV capsid variants, it is now possible to transduce entire neural circuits, rather than individual brain regions. We recently demonstrated that delivery of AAV2.retro expressing a fragment of *HTT* with 85 CAG repeats (AAV2.retro-mHTT85Q) into the adult macaque caudate and putamen results in efficient retrograde transport resulting in mHTT expression and the formation of hallmark inclusion bodies throughout dozens of cortical and sub-cortical striatal afferents (10 weeks post-surgery). In comparison, injection of the parent serotype, AAV2, expressing mHTT85Q resulted in mHTT inclusions primarily within the caudate and putamen, without appreciable retrograde or anterograde transport ^28^. This study laid the groundwork for generating a macaque model of HD that engages the entire cortico-basal ganglia circuit using this new viral vector-mediated approach.

Here, we expanded on these initial efforts and characterized the long-term effects of mHTT85Q expression throughout the macaque cortex and basal ganglia. Following surgery, we assessed cognitive and motor function in a cohort of HTT85Q-treated animals, as well as relevant controls, for a period of 20-months. Additionally, we performed multimodal imaging to assess white and gray matter microstructure, atrophy and patterns of brain-wide functional connectivity. Together, this study describes a new macaque model that closely replicates several of the behavioral and neuropathological changes seen in humans suffering from HD.

## RESULTS

To elicit mHTT85Q expression throughout the caudate and putamen, as well as cortical- and basal ganglia striatal afferents, we took a combinatorial approach and administered a 1:1 mixture of AAV2-HTT85Q and AAV2.retro-HTT85Q at a titer of 2e12 vg/ml (Group 85Q, n=6). Control animals were injected with a 1:1 mixture of AAV2:AAV2retro-HTT10Q (control HTT fragment with 10 CAG repeats; Group 10Q, n=6) or buffered saline only (Group Buffer, n=5), see Table 1. Animals completed a longitudinal battery of cognitive and motor tasks and underwent repeated scans to collect structural (T1w/T2w) magnetic resonance imaging (MRI), diffusion tensor imaging (DTI), and resting state MRI (rsfMRI) data.

**Table 1.** Summary of study participants and surgical cases. Abbreviations: 85Q-fragment of mHTT bearing 85 CAG repeats; 10Q- fragment of mHTT protein bearing 10 CAG repeats, Buffer- phosphate buffered saline. VG- vector genomes, SDR- Spatial Delay Response, DNMS- Delayed Non-Match to Sample. Note that only some of the animals met the training-criteria for the DNMS and 3-choice tasks.

| **ANIMAL**<br>**ID** | **TREATMENT**<br>**GROUP** | **SEX** | **AGE**<br>**AT**<br>**SURGERY**<br>**(YR)** | **WEIGHT**<br>**AT**<br>**SURGERY**<br>**(KG)** | **VECTOR**<br>**DOSE PER**<br>**HEMISPHERE**<br>**(VG)** | **MET**<br>**CRITERIA**<br>**FOR SDR**<br>**TASK** | **MET**<br>**CRITERIA**<br>**FOR DNMS**<br>**TASK** | **MET**<br>**CRITERIA**<br>**FOR**<br>**LIFESAVER**<br>**TASK** |
| --- | --- | --- | --- | --- | --- | --- | --- | --- |
| 1 | 85Q | F | 10.1 | 5.7 | 6.6e11 | Yes | Yes | Yes |
| 2 | 85Q | F | 11.8 | 6 | 6.6e11 | Yes | No | Yes |
| 3 | 85Q | M | 5.9 | 10.7 | 6.6e11 | Yes | Yes | Yes |
| 4 | 85Q | F | 13.0 | 6.2 | 6.6e11 | Yes | Yes | Yes |
| 5 | 85Q | F | 12.3 | 7.5 | 6.6e11 | Yes | Yes | Yes |
| 6 | 85Q | M | 7.1 | 11.1 | 6.6e11 | Yes | Yes | Yes |
| 7 | 10Q | F | 8.7 | 6.5 | 6.6e11 | Yes | No | Yes |
| 8 | 10Q | F | 13.5 | 9.9 | 6.6e11 | Yes | Yes | Yes |
| 9 | 10Q | F | 9.0 | 8.6 | 6.6e11 | Yes | No | Yes |
| 10 | 10Q | M | 7.5 | 8.8 | 6.6e11 | Yes | Yes | Yes |
| 11 | 10Q | F | 11.8 | 6.0 | 6.6e11 | Yes | Yes | Yes |
| 12 | 10Q | M | 10.0 | 9.5 | 6.6e11 | No | No | Yes |
| 13 | Buffer | F | 12.0 | 8.7 | - | Yes | Yes | Yes |
| 14 | Buffer | F | 9.9 | 7.5 | - | No | No | Yes |
| 15 | Buffer | M | 7.7 | 11.3 | - | Yes | Yes | Yes |
| 16 | Buffer | F | 7.9 | 6.0 | - | Yes | Yes | Yes |
| 17 | Buffer | F | 8.8 | 5.9 | - | Yes | No | Yes |

### HTT85Q administration leads to spatial working memory deficits

Disease onset in HD is defined by the presence of disordered hyperkinetic movements, however HD patients also often experience cognitive decline characterized by both spatial and recognition memory impairment, which sometimes precede motor dysfunction ^18, 19, 22, 33, 34^. In NHPs, spatial working memory can be assessed using a 3-choice Spatial Delayed Response (SDR) task, and recognition memory using a Delayed Non-Match to Sample (DNMS) task. Monkeys in this study completed these tasks at baseline as well as 3-, 6-, 9-, 14-, and 20-months post-surgery.

Prior to surgery, 15 animals met the pre-training criteria for the SDR task (**Table 1**), and therefore data from the two remaining subjects were excluded from analysis. For the SDR assay, animals were tasked at remembering the location of a treat-bated, recessed well after a delay of 1, 3, or 5 seconds (**Figure 1A**). A 3-way ANOVA found significant group differences (F(2,12)=17.297, p=0.0003), with no main effect of delay (F(2,24)=0.052, p=0.950) nor timepoint (F(4,48)=1.983, p=0.112). Post-hoc tests revealed that 85Q-treated monkeys performed significantly worse on the SDR task (fewer correct responses) than Buffer- (p=0.0003), and 10Q- (p=0.008) treated monkeys, (**Figure 1B**, delay and timepoint collapsed across groups). Buffer- and 10Q- treated animal performance on the SDR task did not differ significantly (p=0.180). To further explore the time course of the group differences, we conducted planned comparisons between groups at each timepoint. Performance between groups did not differ at the 3-month timepoint; however, significant group differences emerged at the 6-month timepoint and persisted through 20-months, with 85Q animals showing worse performance compared to controls, all p<0.05 (**Figure 1C, Table S1)**.

**Figure 1.**
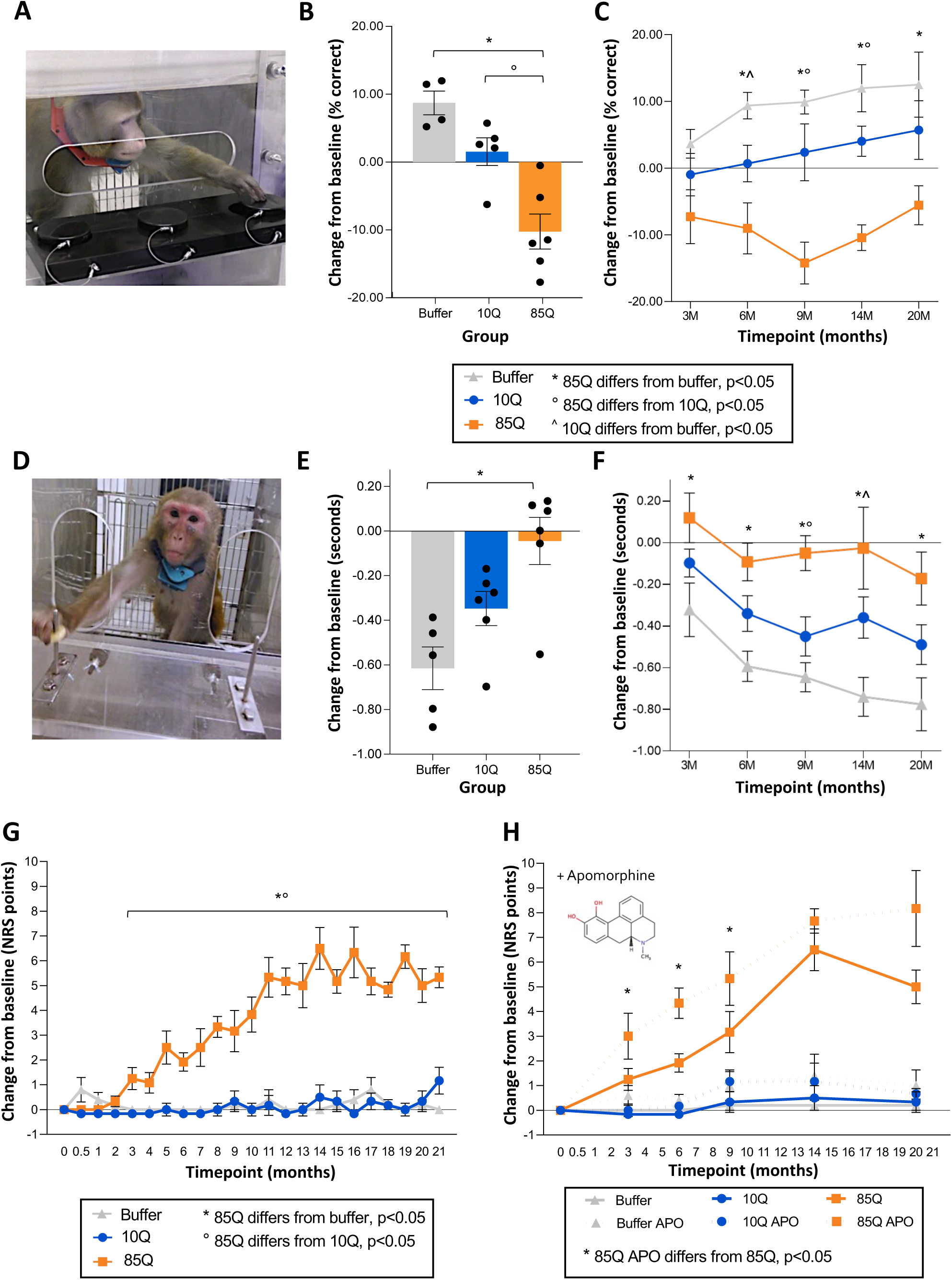
Working memory and motor deficits in 85Q-treated animals. **A)** Example of an animal performing the 3-Choice Spatial Delayed Response Task (SDR). **B)** Change in SDR performance (% correct) on the SDR task, collapsed across timepoints. **C)** Change in SDR performance (% correct), expanded across timepoints. All SDR data are expressed as mean ± SEM (85Q-n=6, 10Q- n=5, Buffer-n=4). The key details significant group differences at each timepoint, demonstrating that working memory changes emerge in this model beginning at 6-months post 85Q administration. **D)** Example of an animal performing the Lifesaver Retrieval Task. **E)** Performance collapsed across timepoint, plotted as the change in Retrieval Latency (seconds) from baseline. **F)** Performance expanded across timepoints, plotted as the change in Retrieval Latency (seconds) from baseline. All Lifesaver Retrieval data are expressed as mean ± SEM (85Q- n=6, 10Q- n=6, Buffer-n=5). The key details significant group differences at each timepoint and show that fine motor skill learning impairment emerges in 85Q animals as early as 3-months post-85Q administration. **G)** Mean changes in motor phenotypes plotted for each group. The key shows significant group differences at each timepoint, indicating that HD motor phenotypes first emerge around 3-months post-surgery. **H)** Neurological Rating Scale (NRS) scores (difference from baseline) pre-apomorphine and post-apomorphine administration plotted for each group, with pre-apomorphine NRS scores depicted by solid lines and post-apomorphine NRS scores depicted by dashed lines. The key depicts significant paired-comparisons at each timepoint, showing that 85Q-treated animals displayed NRS scores that were significantly modulated by apomorphine at 3-9 months post-surgery. All NRS data are expressed as mean ± SEM (85Q- n=6, 10Q- n=6, Buffer-n=5). *p<0.05, 85Q vs. Buffer, °p<0.05, 85Q vs. 10Q, ^^^p<0.05, Buffer vs. 10Q by 3-way ANOVA.

Prior to surgery, 11 animals met the pre-training criteria for the DNMS task, and data from the 6 remaining subjects were excluded from the analysis (**Table 1**). In contrast to the SDR, performance on the DNMS task of recognition memory was unaltered in any of the groups. A 3-way ANOVA confirmed no significant main effects of Group (F(2,8)=0.728, p=0.512), Timepoint (F(4,32)=0.159, p=0.957), nor Delay (F(2,16)=0.007, p=0.993), and no significant interactions between any of these factors. DNMS performance was stable post-surgery across all three groups, suggesting preserved mechanisms of recognition memory in this model.

### HTT85Q administration leads to impaired fine motor skill performance

Pre-symptomatic HD individuals estimated to be 10+ years from disease onset are able to perform self-directed motor tasks (reaching/tapping) similar to age-matched controls, but abnormalities emerge in prodromal HD individuals closer (<5y) to motor diagnosis and persist for many years ^33–35^. To assess fine motor coordination and skill learning, animals completed a Lifesaver Retrieval Task at baseline as well as 3-, 6-, 9-, 14-, and 20-months post-surgery. Prior to surgery, all animals met the pre-training criteria for the Lifesaver Retrieval Task and data from every subject was included in the analysis (Table 1). Treat retrieval latency from a baited metal post was measured separately for each hand (**Figure 1D**). A 3-way ANOVA with repeated measures for the second and third factors (Timepoint, Hand) revealed a significant main effect of Group (F(2,14)=9.035, p=0.003) and Timepoint (F(4,56)=14.774, p<0.0001) on changes in Lifesaver retrieval latencies. The main effect of Hand was not significant (F(1,14)=1.051, p=0.323), therefore, data were subsequently collapsed across this variable. Compared to Buffer controls who became quicker on the task compared to their baseline performance, 85Q- treated animals exhibited significantly longer lifesaver retrieval latencies (p<0.003, **Figure 1E**, data collapsed across timepoints). While 85Q treated animals performed worse on the Lifesaver task compared to 10Q controls, this difference did not reach statistical significance (p=0.101). Performance between Buffer and 10Q-treated controls did not differ significantly (p=0.200). To further explore these effects, we conducted planned group comparisons at each timepoint separately (**Figure 1F** and **Table S2**. 85Q treated animals differed significantly from Buffer controls at all of the timepoints, beginning as early as 3-months post-surgery. 85Q animals and 10Q controls differed significantly at the 9-month timepoint while Buffer and 10Q differed significantly at the 14-month timepoint.

### HTT85Q leads to progressive motor phenotypes that are exacerbated by dopamine modulation

Disease onset in HD is defined by the emergence of motor phenotypes, which are scored using the Unified Huntington Disease Rating Scale (UHDRS) ^36^. To assess similar phenotypes in this model, we developed an NHP-specific neurological rating scale (NRS) based on the motor component of the UHDRS (**Table S3)**. Similar to the UHDRS, NRS rating scale scores range from 0-3, with a score of 0 indicating normal behavior and a score of 3 indicating severely impaired behavior for each phenotype. Monkeys were rated in their homecage at baseline (prior to surgery), 2 weeks post-surgery and then monthly thereafter for 21 months. Scores for each phenotype were summed, resulting in a composite score for each timepoint. A 2-way ANOVA revealed significant effects of Group (F(2,14)=62.709, p<0.0001), and Timepoint (F(21,294)=14.592, p<0.0001), as well as a significant Group x Timepoint interaction (F(42,294)=12.384, p<0.0001) in total NRS scores. Post-hoc tests indicated that 85Q-treated monkeys had significantly higher NRS scores than both Buffer- (p<0.0001) and 10Q- (p<0.0001) treated animals, and that the two control groups did not differ (p=0.999). To further characterize the time course of these effects, we conducted additional planned comparisons between the groups at each timepoint (**Figure 1G, Table S4)**. The results revealed that 85Q had significant increases in their total NRS scores post-surgery compared to Buffer and 10Q beginning at the 3-month timepoint and continuing through the 20-month timepoint. In contrast, comparisons between Buffer and 10Q did not reach significance at any of the timepoints. Refer to the radial plots in **Figure 2** for a breakdown of specific behavioral phenotypes, and their severity, over time. Beginning at 3-months post-surgery, 85Q-treated animals showed infrequent bouts orofacial dyskinesia that included tongue rolling and intermittent tongue protrusion, as well as small changes in forelimb posture, particularly in the distal segment of the limb. As time progressed, 85Q-treated animals exhibited increasingly higher NRS scores, with the emergence of additional postural abnormalities (trunk/hindlimb), slowed treat retrieval, mild forelimb chorea and slowed homecage ambulation. By 9 months post-surgery, animals showed mild evidence of hindlimb bradykinesia that worsened over time.

**Figure 2.**
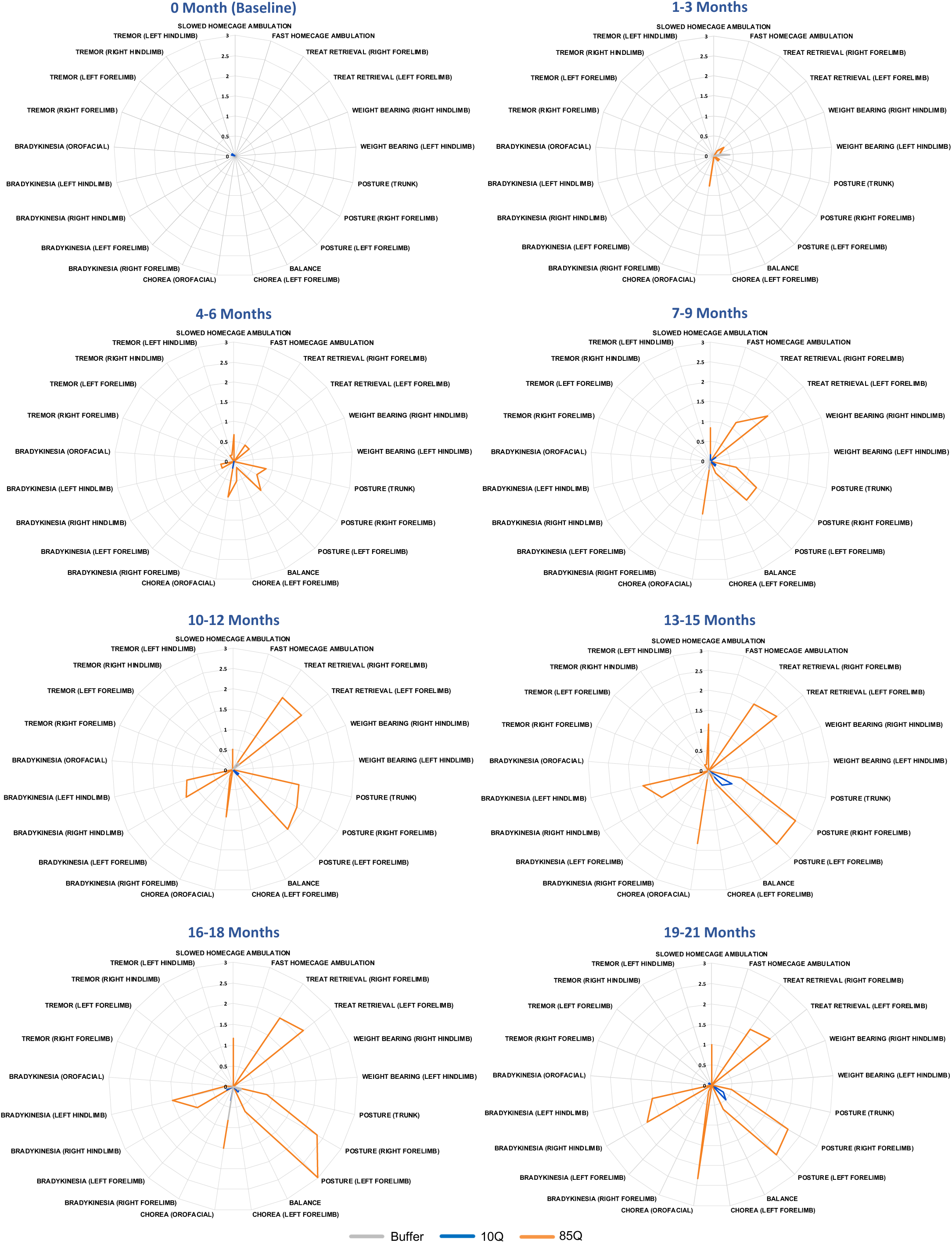
Motor phenotypes characterized using an NHP-specific Neurological Rating Scale. Scores from individual NRS categories were collapsed into 3-month time bins and plotted on radar charts. Early developing phenotypes of 85Q-treted animals included mild orofacial chorea, impairments in treat retrieval and aberrant forelimb posture that were observed in the first 3 months. By 6 months post-surgery, 85Q treated animals had additional postural abnormalities (trunk/hindlimb), slowed homecage ambulation, mild hindlimb tremor and some instances of hindlimb bradykinesia. Each of these phenotypes persisted and worsened (higher NRS scores) over time out to 21 months post-surgery. Some, but not all, buffer and 10Q control treated animals showed evidence of mild motor phenotypes.

Motor phenotypes exhibited by HD patients are modulated by the neurotransmitter, dopamine. Dopamine receptor agonists have been shown to exacerbate abnormal involuntary movements in early HD patients ^37^ and current FDA-approved drugs to treat chorea in HD are the vesicular monoamine transporter 2 (VMAT2) inhibitors, Tetrabenazine^38^ and Deutetrabenazine^39^, which reduce dopamine availability in the synapse. In order to probe the involvement of dopamine system dysfunction in our model, we assessed the impact of the non-selective dopamine receptor agonist, apomorphine, on motor behavior. Animals were scored using the NRS immediately prior to, as well as directly following, intramuscular apomorphine administration (0.3 mg/kg) at baseline and at 3-, 6-, 9-, 14- and 20-months post-surgery. NRS scores from 85Q-treated animals were significantly exacerbated by apomorphine, compared to controls. A 3-way ANOVA confirmed this observation, revealing significant main effects of Group (F(2,14)=62.441, p<0.0001), Timepoint (F(4,56)=11.807, p<0.0001), and ApoCondition (F(1,14)=22.552, p<0.0001), as well as significant interaction effects between Timepoint and Group (F(8,56)=5.967, p<0.0001), and between ApoCondition and Group (F(2,14)=6.071, p=0.013). To further characterize these effects, we conducted additional planned comparisons between pre- and post-Apomorphine scores at each time point for each group separately (**Figure 1H, Table S5)**. Results indicated that 85Q -treated animals had significantly elevated NRS scores following apomorphine administration, compared to pre-apomorphine administration, at the 3-, 6-, and 9-month timepoints, but not at the 14- nor 20-month timepoints. In contrast, changes in pre- vs post-Apomorphine administration scores did not significantly differ at any of the timepoints for Buffer- or 10Q- treated controls.

### mHTT delivery results in microstructural changes in several white matter fiber tracts

White matter (WM) fiber tracts can be imaged using diffusion tensor imaging (DTI), with the most commonly reported metric being fractional anisotropy (FA) in water diffusion. Reduced FA is observed in HD in many WM regions including the corpus callosum, superior and inferior longitudinal fasciculi, corona radiata, internal capsule and cingulum, indicating microstructural changes to the axon including reduced axonal integrity and demyelination ^11–14^. To query white matter changes in our model, DTI data were collected pre-surgery and 3-, 6-, 9-, 14-, and 20-months post-surgery. Parameter maps of FA, axial diffusivity (AD), radial diffusivity (RD) and mean diffusivity (MD) were generated in ONPRC18 template space ^40^. **Figure 3A** illustrates the results of voxel-wise comparisons of FA, using a WM mask, conducted between the baseline (pre-surgical) timepoint and subsequent timepoints for each group separately. As early as 3-months post-surgery, 85Q-treated monkeys had several regions of WM with significant (p<0.01) reductions in FA that persisted through the 20-month timepoint; in contrast, no changes in FA were observed in 10Q- and Buffer- treated animals.

**Figure 3.**
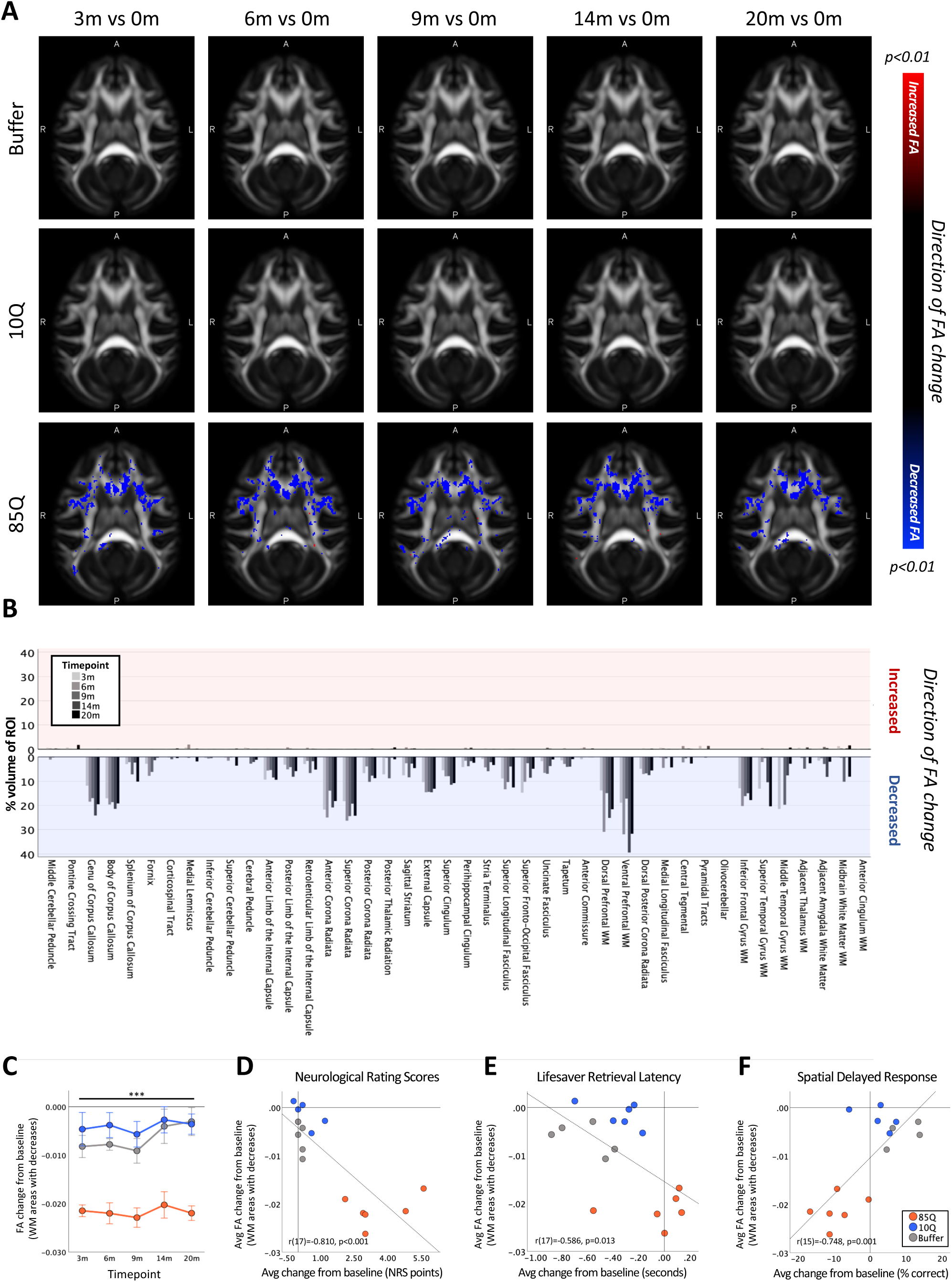
85Q-mediated microstructural changes in cerebral white matter. **A)** ONPRC18 FA template with overlaying p-value maps shown at a threshold of p<0.01. Blue voxels indicate regions of significant FA decrease, and red voxels indicate regions of significant FA increase. Although there were slight changes in FA in the Buffer and 10Q treated animals over time, none of the contrasts reached statistical significance. **B)** Histogram illustrating the percent volume of each ROI where significant FA changes were identified. Top panel shows increased FA (corresponding to the red voxels in 3A), bottom panel shows decreased FA (corresponding to the blue voxels in panel **A**). A mask that merged together the thresholded p-value maps from each timepoint was created and the line chart in (**C**) illustrates the average magnitude of change in FA under the mask for each group separately. Data are expressed as mean ± SEM (85Q- n=6, 10Q- n=6, Buffer-n=5), Repeated Measures ANOVA. Scatterplots illustrating two-tailed Pearson correlations between FA decreases in WM and behavior (both collapsed across timepoint) for three different behavioral measures; **D**) NRS, **E**) Lifesaver Retrieval Task and **F**) SDR Task. FA- fractional anisotropy, ROI- region of interest. ***p<0.001; 85Q differs from Buffer and 10Q.

To characterize the distribution of the significant voxel-wise differences, WM labels from the ONPRC18 macaque brain atlas were applied to the thresholded p-value maps from 85Q-treated animals, and the percent volume of significant FA reduction was calculated for each WM ROI at each timepoint **(Figure 3B)**. Broadly, there was an anterior-posterior spatial distribution of WM changes in 85Q animals, with larger relative volumes of reduced FA occurring in more anterior ROIs, without robust hemispheric differences. Of particular note was the large percentages of the dorsal and ventral prefrontal WM tracts, as well as the anterior/dorsal corona radiata, internal capsule, external capsule, and the genu and body of the corpus callosum exhibiting significant reductions in FA compared to baseline. To characterize the overall time-course of these changes, average change in FA (from baseline) in significant regions were calculated for each animal at each post-surgical timepoint and plotted in **Figure 3C**. A repeated measure ANOVA revealed significant main effects of Group (F(2,14)=70.909, p<0.0001), and Timepoint (F(4,56)=2.710, p=0.039), but no interaction between these factors (F(8,56)=0.618, p=0.759). Post-hoc tests revealed that 85Q-treated animals had significantly greater decreases in FA after surgery compared to Buffer (p<0.0001), and 10Q (p<0.0001) animals, whereas Buffer- and 10Q-treated animals did not differ significantly in the magnitude of FA changes (p=0.052). To query the relationship between the observed DTI changes and behavioral impairments, we computed correlations between FA changes and behavioral scores that were collapsed across all 5 post-surgical timepoints. FA decreases in WM were correlated with all three behavioral measures, such that greater decreases in FA were significantly associated with higher NRS scores (r(17)=-0.810,p<0.001) (**Figure 3D)**, longer Lifesaver retrieval latencies (r(17)=-0.586, p=0.013) (**Figure 3E)** and lower percentages of correct responses on the 3-Choice SDR task (r(15)=-0.748, p=0.001) (**Figure 3F)**.

Additional analysis of DTI parameters including radial diffusivity (RD), mean diffusivity (MD) and axial diffusivity (AD) were performed, following the strategy described for FA. 85Q-treated animals showed significant elevations in RD (**Figure S1**) and MD (**Figure S2**), as well as significant decreases in AD (**Figure S3**), compared to controls (p<0.01 for each measurement). These changes followed similar spatiotemporal trajectories to the changes observed in FA. In particular, there were areas of significant RD and MD increase in corresponding regions of FA decrease in the dorsal and ventral prefrontal WM tracts, as well as areas of the anterior/dorsal corona radiata, internal capsule, external capsule, and the genu and body of the corpus callosum (**Figure S1-2**). Brain-wide, the spatial distributions of RD and MD increases generally overlapped the FA decreases, whereas the significant decreases in AD were more widely distributed **(Figure S4)**. None of the changes in AD, RD or MD observed in the 10Q and Buffer groups survived thresholding at the p<0.01 level.

#### Altered diffusion in cortex and basal ganglia

More recently, a pattern of increased FA in the putamen and globus pallidus has been described in pre-manifest/early symptomatic HD patients; and a pattern of decreased FA in gray matter in areas including the ventromedial frontal cortex, insula cortex, and the cerebellum has been described in advanced HD patients ^11, 41–44^. To explore whether similar changes in FA are evident in gray matter (GM) regions in our HD macaques, we conducted voxel-wise comparisons between the baseline (pre-surgical) FA maps and subsequent timepoints for each group separately using a GM mask. By 3-months post-surgery, 85Q-injected animals had areas of the caudate as well as large regions of cortex with significant reductions in FA, and areas of the putamen and globus pallidus with significant increases in FA (p<0.01 threshold, **Figure 4A**). In contrast, none of the changes observed in the 10Q- and Buffer-injected animals survived thresholding at the p<0.01 level.

**Figure 4.**
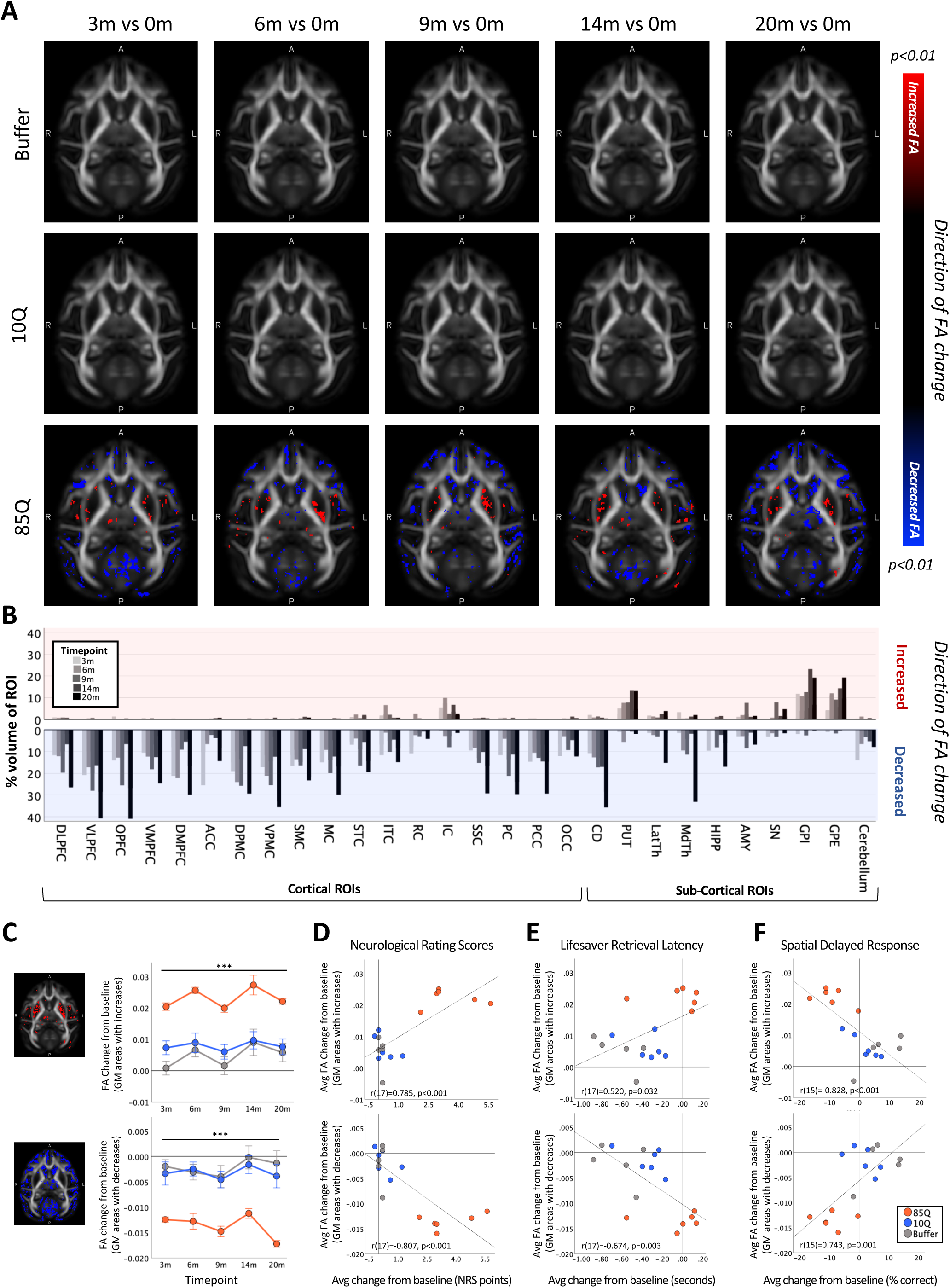
85Q-mediated microstructural alterations in cortico-basal ganglia grey matter. **A)** ONPRC18 FA template with overlaying p-value maps shown at a threshold of p<0.01. Red voxels indicate regions of significant FA increase, and blue voxels indicate regions of significant FA decrease. **B)** Histogram illustrating the percent volume of each cortical and subcortical ROI where significant FA changes were identified. Top panel shows increased FA (corresponding to the red voxels in **A**), bottom panel shows decreased FA (corresponding to the blue voxels in **A**). **C)** Masks that merged together the thresholded p-value maps from each timepoint were created. Line charts illustrate the average magnitude of FA changes under each mask for each group separately. Data are expressed as mean ± SEM (85Q- n=6, 10Q- n=6, Buffer-n=5), Repeated Measures ANOVA. Scatterplots illustrating two-tailed Pearson correlations between FA changes in GM and behavior (both collapsed across timepoint) for three different behavioral measures; **D**) NRS, **E**) Lifesaver Retrieval Task and **F**) SDR Task. FA- fractional anisotropy, ROI- region of interest. ***p<0.001; 85Q differs from Buffer and 10Q. *Abbreviations: DLPFC, dorsolateral prefrontal cortex; VLPFC, ventrolateral prefrontal cortex; OPFC, orbitofrontal cortex; VMPFC, ventromedial prefrontal cortex; DMPFC, dorsomedial prefrontal cortex; ACC, anterior cingulate cortex; DPMC, dorsal premotor cortex; VPMC, ventral premotor cortex; SMC, supplemental motor cortex; MC, primary motor cortex; STC, superior temporal cortex; ITC, inferior temporal cortex; RC, rhinal cortex; IC, insular cortex; SSC, somatosensory cortex; PC, parietal cortex; PCC, posterior cingulate cortex; OCC, occipital cortex; CD, caudate; PUT, putamen; LatTH, lateral thalamus; MdTH, medial thalamus; HIPP, hippocampus; AMY, amygdala; SN, substantia nigra; GPI, internal globus pallidus; GPE, external globus pallidus; LatVentricle, lateral ventricles*.

GM labels in the ONPRC18 atlas were applied to the thresholded p-value maps from 85Q-treated animals, and the percent volume of significant increases and reductions in FA were calculated for each ROI at each timepoint **(Figure 4B)**. Numerous cortical and subcortical brain regions showed reduced FA; while the majority of the regions of increased FA were located in subcortical areas. Of particular note was the divergent pattern of significant changes in the striatum/basal ganglia in 85Q-treated animals, with regions of the caudate showing significant FA decreases and regions of the putamen and globus pallidus showing significant FA increases compared to baseline.

To characterize the overall time-course of these changes, masks that merged together the thresholded p-value maps from each timepoint were created separately for regions of FA increase and regions of FA decrease. The change in FA (from baseline) under the masks was calculated for each animal at each post-surgical timepoint and plotted in **Figure 4C**. In GM regions with significantly increased FA, a 2-way repeated measure ANOVA revealed significant main effects of Group (F(2,14)=32.190, p<0.0001), and Timepoint (F(4,56)=6.288, p=0.0003), but no interaction between these factors (F(8,56)=0.560, p=0.806). Post-hoc tests revealed that the 85Q group had significantly greater increases in FA after surgery compared to the Buffer group (p<0.0001), and the 10Q group (p<0.0001), whereas the Buffer and 10Q groups did not differ significantly (p=0.999). Similarly, in GM regions with significantly reduced FA, a 2-way repeated measure ANOVA revealed a significant Group by Timepoint interaction (F(8,56)=2.804, p=0.011) as well as significant main effects of Group (F(2,14)=34.080, p<0.0001), and Timepoint (F(4,56)=8.690, p<0.0001). Post-hoc tests indicated that 85Q-treated monkeys had significantly greater decreases in GM FA after surgery compared to Buffer- (p<0.0001), and 10Q- (p<0.0001) treated monkeys, whereas Buffer and 10Q did not differ significantly (p=0.999).

To probe the relationship between the observed DTI changes in GM and the behavioral measures, two-tailed Pearson correlations were computed between FA and behavioral change scores that had been collapsed across all 5 post-surgical timepoints. FA changes in GM were correlated with all three behavioral measures, such that greater increases in FA were significantly associated with higher NRS scores (r(17)=0.785,p<0.001), **(Figure 4D, top row),** longer Lifesaver retrieval latencies (r(17)=0.520, p=0.032) (**Figure 4E, top row**) and lower percentage correct on the 3-Choice SDR task (r(15)=-0.828, p<0.001) (**Figure 4F, top row**). Additionally, greater decreases in FA were significantly associated with higher NRS scores (r(17)=-0.807,p<0.001) **(Figure 4D, bottom row)**, longer Lifesaver retrieval latencies (r(17)=-0.674, p=0.003) **(Figure 4E, bottom row)** and lower percentage correct on the 3-Choice SDR task (r(15)=0.743, p=0.001) **(Figure 4F, bottom row).**

#### Cortico-striatal atrophy revealed with Tensor Based Morphometry

Tensor Based Morphometry (TBM) is a technique used to assess changes in brain morphometry by comparing deformation fields that align anatomical MR images to a template image. Applied longitudinally, changes in these deformation fields can be used to characterize 3D patterns of structural change. Atrophy of many brain regions (caudate, putamen, thalamus, multiple regions of cortex) has been described in symptomatic HD patients using this technique ^45, 46^, and recent longitudinal data indicate that some changes begin prior to disease onset; particularly in regions of the putamen and globus pallidus ^12, 45, 47^. High-resolution T2-weighted SPACE scans were collected pre-surgery and 3-, 6-, 9-, 14-, and 20-months post-surgery. For each timepoint, deformation fields aligning with the ONPRC18 template were computed, and log Jacobian Determinant maps, reflecting local structural changes, were generated. In the resulting maps, positive values indicated areas where individual space was dilated to align with ONPRC18 space (and was therefore smaller), whereas negative values indicate areas where individual space was contracted to align with ONPRC18 space (and was therefore larger). Using a GM mask, voxel-wise comparisons were conducted to compare the baseline (pre-surgical) timepoint and subsequent timepoints. As early as 3-months post-surgery, 85Q-treated monkeys exhibited regions of the basal ganglia, prefrontal- and premotor-cortex with significant tissue contractions (larger log Jacobian Determinants), indicating tissue atrophy, compared to baseline that persisted through the 20-month timepoint (p<0.01). In contrast, the none of the changes observed in 10Q and Buffer survived thresholding at the p<0.01 level **(Figure 5A)**. GM labels available in the ONPRC18 atlas were applied to the thresholded p-value maps from 85Q-treated animals, and the percent volumes of significant log Jacobian Determinant increases were calculated for each ROI at each timepoint. Major brain areas showing tissue contraction were the caudate, putamen, globus pallidus (internal and external segments), as well as numerous frontal and motor cortical areas, with no hemispheric differences (**Figure 5B)**.

**Figure 5.**
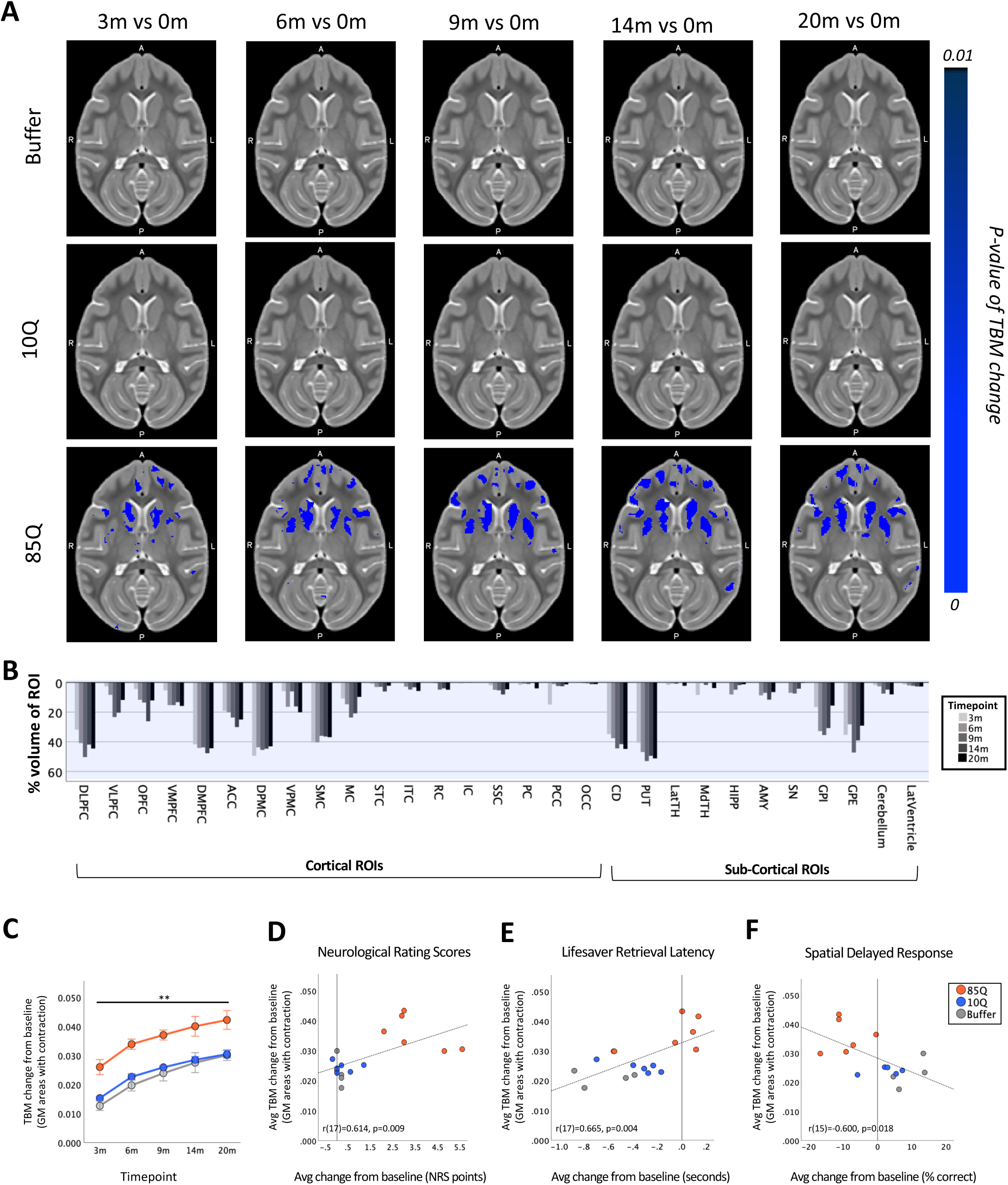
85Q-mediated tissue atrophy in cortico-basal ganglia grey matter. **A)** ONPRC18 T2w template with overlaying p-value maps shown at a threshold of p<0.01. Blue voxels indicate regions of significant TBM contraction (increased log Jacobian Determinants). **B)** Histogram illustrating the percent volume of each cortical and subcortical ROI where significant TBM changes were identified (corresponding to the blue voxels in **A**). **C)** A mask that merged together the thresholded p-value maps from each timepoint was created. Line charts illustrate the average magnitude of log Jacobian Determinate changes under this mask for each group separately. Data are expressed as mean ± SEM (85Q- n=6, 10Q- n=6, Buffer-n=5), Repeated Measures ANOVA. **D)** Scatterplots illustrating two-tailed Pearson correlations between TBM changes in GM and behavior (both collapsed across timepoint) for three different behavioral measures; **D**) NRS, **E**) Lifesaver Retrieval Task and **F**) SDR Task. TBM- tensor based morphometry, ROI- region of interest. **p<0.01; 85Q differs from Buffer and 10Q. *Abbreviations: DLPFC, dorsolateral prefrontal cortex; VLPFC, ventrolateral prefrontal cortex; OPFC, orbitofrontal cortex; VMPFC, ventromedial prefrontal cortex; DMPFC, dorsomedial prefrontal cortex; ACC, anterior cingulate cortex; DPMC, dorsal premotor cortex; VPMC, ventral premotor cortex; SMC, supplemental motor cortex; MC, primary motor cortex; STC, superior temporal cortex; ITC, inferior temporal cortex; RC, rhinal cortex; IC, insular cortex; SSC, somatosensory cortex; PC, parietal cortex; PCC, posterior cingulate cortex; OCC, occipital cortex; CD, caudate; PUT, putamen; LatTH, lateral thalamus; MdTH, medial thalamus; HIPP, hippocampus; AMY, amygdala; SN, substantia nigra; GPI, internal globus pallidus; GPE, external globus pallidus; LatVentricle, lateral ventricles*.

To characterize the overall time-course of these changes, masks that merged together the thresholded p-value maps from each timepoint were created for areas of significant contraction. The change in the log Jacobian Determinant maps (from baseline) under the masks was then calculated for each animal at each post-surgical timepoint and plotted in **Figure 5C**. In the regions with significant increases in log Jacobian Determinants, a 2-way repeated measure ANOVA revealed significant main effects of Group (F(2,14)=5.384, p=0.0003), and Timepoint (F(4,56)=89.068, p<0.0001), but no interaction between these factors (F(8,56)=0.234, p=0.983). Post-hoc tests indicated that 85Q-treated animals had significantly greater tissue contractions (greater increases in average log Jacobian Determinant values) compared to Buffer- (p=0.003), and 10Q- (p=0.002) treated animals, with no differences between control groups (p=0.999).

Similar to other imaging measurements, correlations were characterized between the observed TBM results and the behavioral measures; two-tailed Pearson correlations were computed between log Jacobian Determinant changes and behavioral change scores that had been collapsed across all 5 post-surgical timepoints. In regions with significant tissue atrophy, TBM changes in GM were correlated with all three behavioral measures, such that greater contractions were significantly associated with higher NRS scores (r(17)=0.614, p=0.009), **(Figure 5D)**, longer Lifesaver retrieval latencies (r(17)=0.665, p=0.004) (**Figure 5E**) and lower percentages correct on the 3-Choice SDR task (r(15)=-0.600, p=0.018) (**Figure 5F**).

#### Reduced cortico-basal ganglia functional connectivity

Resting-state fMRI (rsfMRI) is a technique used to identify areas of the brain that exhibit correlated fluctuations in blood oxygenation level dependent (BOLD) MR signal intensity in the absence of a specific stimulus (hence, in the “resting state”) ^48^. Independent component analysis (ICA) of rsfMRI is a data-driven computational method used to decompose correlated patterns of BOLD signals into networks (or “components”) with distinct spatial and temporal characteristics; without the reliance on pre-established user-defined ROIs for seed-based correlations. Using ICA, we identified 4 independent resting-state components (ICs) of interest using data collected at the baseline timepoint from all of the study animals (**Figure S5-S6)**. IC1 includes areas of the occipital cortex and ventrolateral PFC; IC2 includes the caudate, putamen, ventral sensory-motor cortex and medial PFC; IC3 includes the caudate and several prefrontal and temporal cortical regions; and IC4 includes the caudate, putamen, cingulate, dorsal-prefrontal and dorsal-motor areas (**Figure S5**). To facilitate longitudinal group-level comparisons, dual-regression (DR) analysis was applied to each timepoint, focusing on these 4 ICs, and z-score connectivity maps for each of the 4 ICs from each monkey at each timepoint were subsequently generated.

In the field of HD, studies applying rsfMRI with ICA-DR ^49–52^ indicate that areas of decreased RSFC emerge first in pre-manifest HD-gene carriers, which shifts later in disease stages to increased RSFC, potentially as a compensatory mechanism. To define changes in RSFC in our model, we conducted voxel-wise comparisons for each IC between z-score maps from the baseline (pre-surgical) timepoint and subsequent timepoints using a whole brain mask. The most robust differences were observed in the component that overlapped striatal and sensory-motor cortex (IC2) (**Figure 6, Figure S5**). As early as 3-months post-surgery and continuing through the 20-month timepoint, 85Q-treated monkeys showed significant decreases in IC2 z-scores in many cortical and subcortical areas (p<0.01). In contrast, none of the changes observed in 10Q and Buffer survived thresholding at the p<0.01 level **(Figure 6A)**. The percent areas of significant change in z-scores with IC2 were calculated for each ROI in the ONPRC18 labelmap at each timepoint and plotted in **Figure 6B**. The results highlighted regions of the VLPFC, VMPFC, ITC, IC as well as areas of the putamen, substantia nigra, globus pallidus, amygdala, hippocampus, and cerebellum with significantly reduced connectivity (p<0.01) to the IC2 network over the 20-month study timeline.

**Figure 6.**
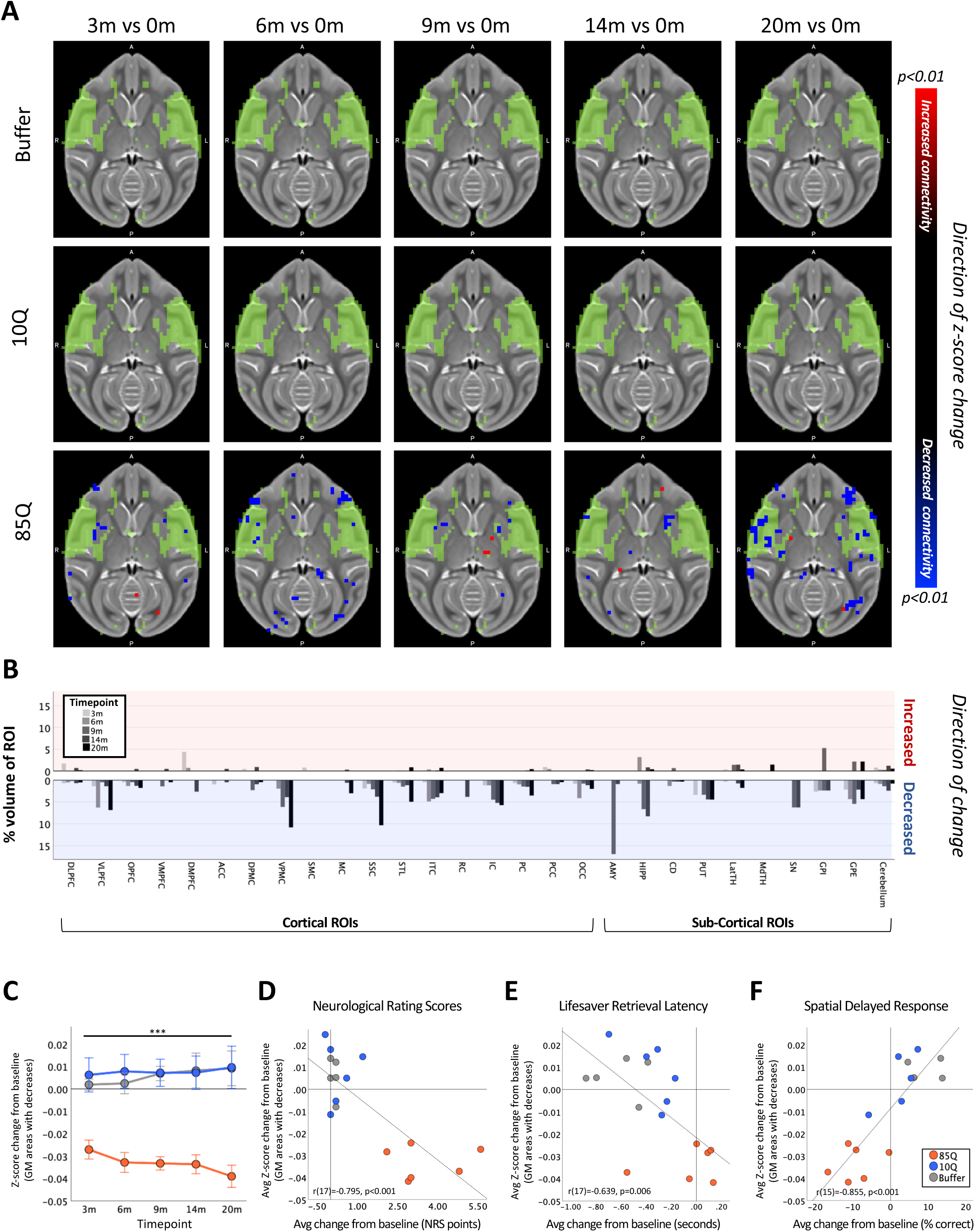
85Q-mediated alterations in patterns of brain-wide resting-state functional connectivity. **A)** ONPRC18 T2w template with overlaying map of Independent Component 2 (IC2), shown in green. Additional overlaying p-value maps are shown at a threshold of p<0.01. Blue voxels indicate regions of significantly reduced RSFC (decreased z-score) with IC2, and red voxels indicate regions of significant increased RSFC (increased z-score) with IC2. **B)** Histogram illustrating the percent volume of each cortical and subcortical ROI where RSFC changes were identified, extracted using the GM parcellations available in the ONPRC18 atlas. Top panel shows increased z-scores (corresponding to the red voxels in A), bottom panel shows decreased z-scores (corresponding to the blue voxels in **A**). **C)** A mask that merged together the thresholded p-value maps from each timepoint was created for regions of RSFC decreases with IC2. Line charts illustrate the average magnitude of RSFC changes (from baseline) under this mask for each group separately, +/-1 SEM. **D)** Scatterplots illustrating two-tailed Pearson correlations between RFSC changes in GM and behavior (both collapsed across timepoint) for three different behavioral measures; **D**) NRS, **E**) Lifesaver Retrieval Task and **F**) SDR Task. RSFC- resting state functional connectivity. ***p<0.001; 85Q differs from Buffer and 10Q. *Abbreviations: DLPFC, dorsolateral prefrontal cortex; VLPFC, ventrolateral prefrontal cortex; OPFC, orbitofrontal cortex; VMPFC, ventromedial prefrontal cortex; DMPFC, dorsomedial prefrontal cortex; ACC, anterior cingulate cortex; DPMC, dorsal premotor cortex; VPMC, ventral premotor cortex; SMC, supplemental motor cortex; MC, primary motor cortex; STC, superior temporal cortex; ITC, inferior temporal cortex; RC, rhinal cortex; IC, insular cortex; SSC, somatosensory cortex; PC, parietal cortex; PCC, posterior cingulate cortex; OCC, occipital cortex; CD, caudate; PUT, putamen; LatTH, lateral thalamus; MdTH, medial thalamus; HIPP, hippocampus; AMY, amygdala; SN, substantia nigra; GPI, internal globus pallidus; GPE, external globus pallidus; LatVentricle, lateral ventricles*.

To characterize the overall time-course of these changes, the change in IC2 z-scores (from baseline) in these significant regions were calculated for each animal at each post-surgical timepoint and plotted in **Figure 6C**. A 2-way repeated measure ANOVA revealed a significant main effect of Group (F(2,14)=27.716, p<0.0001), but no effect of Timepoint (F(4,56)=0.043, p=0.996) nor interaction (F(8,56)=0.678, p=0.708). Post hoc tests indicated that 85Q animals had significantly larger decreases in IC2 z-scores after surgery compared to the Buffer (p<0.0001), and 10Q (p<0.0001) animals, with no significant differences in the magnitude of z-score changes between controls (p=0.989).

To probe the relationship between the changes in IC2 connectivity and the behavioral measures, correlations were computed between the z-score changes and behavioral change scores, collapsed across all 5 post-surgical timepoints. Similar to the changes we observed using the other imaging modalities, the magnitude of decreases in RSFC with IC2 correlated with all three behavioral measures such that larger decreases in RSFC were associated with higher NRS scores (r(17)=-0.795, p<0.001) (**Figure 6D**), longer Lifesaver retrieval latencies (r(17)=-0.639, p=0.006) (**Figure 6E**) and lower percent correct on the 3-Choice SDR task (r(15)=0.855, p<0.001) (**Figure 6F**).

There were also modest, and transient changes in the functional-connectivity of IC1, IC3 and IC4 over the 20-month study timeline. Additional analysis indicated that in 85Q-treated monkeys, areas of the occipital and prefrontal cortex exhibited an initial increase in connectivity with IC1 that returned to baseline levels by 6-months post-surgery (**Figure S7**). The changes observed in IC3 and IC4 were more subtle (**Figure S8-9**), showing only small regions of cortex with altered RSFC compared to baseline. In all cases, however, none of the changes in IC1, IC3 or IC4 observed in 10Q and Buffer survived thresholding at the p<0.01 level.

## DISCUSSION

The goal of this study was to develop a macaque model of HD that recapitulates the progressive sequelae of neurodegenerative changes throughout the cortico-basal ganglia network and gives rise to characteristic motor and cognitive decline seen in human HD patients. Secondarily, we sought to characterize the potential relationships between behavioral phenotypes and neuroimaging findings of disease progression in order to define a set of reliable outcome measures that can be used in future translational studies of candidate HD therapeutics in this model. Because the striatum is the first brain region affected in HD, with degeneration extending to white matter and eventually widespread regions of the cortex^53^, we undertook a unique approach delivering a mixture of AAV2 and AAV2.retro in order to express mHTT throughout the entire macaque cortico-striatal circuit.

Over the 20-month study, HTT85Q animals developed motor and cognitive deficits that were accompanied by microstructural alterations, atrophy, and reduced functional connectivity throughout the cortico-basal ganglia network. Due to the long scanning times achievable with anesthetized NHPs, high-resolution DTI enabled a more detailed analyses of white matter tracts, in addition to cortical and sub-cortical gray matter brain structures than has been previously achieved. The reduction in white matter FA, a biomarker associated with myelin breakdown and axonal swelling, seen in HTT85Q macaques closely mirrors changes seen in human HD patient white matter tracts, including the prefrontal WM tracts, corona radiata, internal and external capsules, and the corpus callosum^43^. Interestingly, the caudate and several widespread cortical regions also exhibited reduced FA as well, a phenomenon less reported in human HD cases, and which could potentially indicate atrophy affecting dendritic processes in these regions. In contrast, areas of the putamen and globus pallidus exhibited significantly increased FA in HTT85Q-treated animals, which has also been described in both pre-manifest and symptomatic HD patients ^41–44^ and speculated to result from the of loss of radially-dispersed fibers projecting from striatal medium spiny neurons ^44^. We hypothesize that increased FA in the putamen and globus pallidus may have also resulted secondarily from an increase in gliotic processes, affecting water diffusion in these areas of degeneration, and histological studies are planned when brain tissue from these animals becomes available. In addition to microstructural changes detected by DTI, HTT85Q-treated animals also exhibited subtle striatal and cortical atrophy, along with mild reductions in network connectivity, which were present by the first time point of analysis (3-months post AAV-administration) (Figure S10). The overall pattern of neurodegeneration that we characterized in this study models similar changes detected in prodromal and early-stage HD patients ^12, 45, 47, 49–52^. Future evaluation at even earlier timepoints will help address whether degenerative processes begin in the striatum first, before being detectable in white matter and afferent cortical areas-as is seen in human HD cases- or whether these pathological changes occur simultaneously in this AAV-based model. HTT85Q-treated animals developed progressive deficits in fine motor skill learning and spatial working memory that emerged at 3- and 6-months post-AAV administration, respectively. However, they maintained their ability to recognize novel objects, similar to data reported from the PREDICT-HD study indicating that spatial working memory appears to be more impacted in premanifest/early-stage HD than recognition memory ^33^. This finding may also result from the specific pattern of mHTT expression in this model, as we previously showed that AAV2.retro delivery of HTT85Q to the caudate and putamen results in higher expression of mHTT protein and aggregate expression in prefrontal cortical areas and the caudate, regions critical for spatial working memory performance, compared to the hippocampus and medial temporal cortices which play a more substantial role in recognition memory.^28^ Moreover, recruitment of the medial temporal lobe in recognition memory is even more substantial when large, novel picture libraries are used as stimuli, as was employed here, versus highly familiar picture libraries that generate proactive interference.^54^

Beginning at 3 months post-surgery, 85Q-treated macaques also developed mild motor phenotypes similar to those experienced by HD patients including incoordination, forelimb and orofacial chorea, hindlimb tremor and postural changes in the distal portion of the forelimbs. Over time, these behaviors worsened in severity and further changes were observed including evidence of delayed gait initiation and hindlimb bradykinesia during locomotion (Figure 2). Interestingly, the severity of these phenotypes was exacerbated by the dopamine agonist, apomorphine, at early timepoints (3-9 months post-surgery), but was not maintained at the 14-and 20-month timepoints. Apomorphine has a high binding affinity for dopamine D_2_, D_3_ and D_5_ receptors and we postulate that the lack of behavioral exacerbation at the latter timepoints may have resulted from a progressive reduction in dopamine receptors in the basal ganglia, resulting from mHTT expression. Altered dopamine neurotransmission has been well documented in HD; for review, see Cepeda et al ^55^. Positron emission tomography (PET) studies have confirmed a progressive loss of striatal D2 and D1 receptor density throughout the course of disease that correlate with clinical severity assessed using motor UHDRS and Total Functional Capacity (TFC) scales. ^56, 57^ PET studies are planned to characterize potential dopamine receptor density alterations in this model. Overall, similar to the degree of imaging findings, the cognitive and motor phenotypes exhibited by the HTT85Q macaques appear to model symptoms experienced by patients in the early stages of their disease development (**Figure S10**).

While the mouse models of HD developed to date have clear advantages (e.g., the relative ease of generating transgenic and knock-in animals, and the fact that they model several of the mHTT-mediated neuropathological intracellular cascades seen in human HD), they do not replicate the complex array of behavioral symptoms seen in human patients, including chorea. Moreover, their much smaller brain size (∼3700- fold smaller by weight) makes scaling up drug delivery strategies to humans an incredibly arduous task, particularly with a delivery that requires complex neurosurgery to deep brain structures. Work with large animal models of HD has accelerated greatly over the past decade due to the development of viral-based and genetically modified sheep, minipigs and NHPs ^27, 29, 58, 59^ For a comprehensive review, see Howland et al, 2020. ^26^ Transgenic HD sheep do not exhibit many of the overt behavioral phenotypes seen in human HD patients, aside from circadian disturbances, but do show signatures of mHTT-mediated pathology including the development of cortical mHTT-positive inclusions, evidence of increased brain urea, as well as loss of cannabinoid receptor 1 (CB1) and dopamine-and cyclic AMP-regulated phosphoprotein (DARPP-32) in the globus pallidus.^58, 60–62^ While HD sheep do not develop cognitive and motor phenotypes, at least out to 5 years of age, an advantage of this model over existing transgenic and viral-mediated minipig and NHP HD models is that it was created using cDNA for full length human *mHTT (*67 exons), providing a full landscape for HTT-lowering therapeutic constructs^59^. Transgenic HD minipigs also have a very long prodromal period, show evidence of mHTT-positive inclusions in striatum and cortex, reduced DARPP-32 expression, microgliosis and mild white matter demyelination only beginning at 2 years of age.^63^ At 4-6 years of age, HD minipigs show more profound neuropathological changes and only at 6-8 years of age do they develop a mild motor phenotype, including impaired gait and reduced treat retrieval by tongue. More recently, a knock-in minipig model has been created by Exemplar Genetics that expresses full-length mHTT-150Q in all tissues evaluated, and piglets show uncoordinated hindlimb movement (unpublished data). Further characterization in this model is ongoing. The sheep and pig models have been employed in pre-clinical HD research to screen promising HTT-lowering therapeutics and one of these approaches, AAV-mediated delivery of a mHTT-specific microRNA, has recently advanced to the clinical trial stage (AMT-130), illustrating just how critical these large animal models are to the field^64, 65^.

In comparison to the transgenic HD sheep and minipigs, the HD NHPs reported here have a much shorter premanifest period with mild motor phenotypes already present by 3 months post-surgery, and working memory reductions at 6-months post-surgery. Previous work by our group shows that mHTT inclusions are present throughout the cortex and basal ganglia in this model by 10-weeks post-surgery.^28^ Additionally, neuroimaging outcome measures were significantly altered by 3 months post-surgery. In addition to this much shorter prodromal period, AAV2.retro-mediated HD macaques display many of the cardinal features of HD that are used as primary and secondary outcome measures in HD human clinical trials including motor and cognitive decline measured via the UHDRS and TFC (and via the NHP clinical rating scale in this model) and regional brain atrophy that is quantified using MRI/DTI (**Figure S10**). Importantly, the behavioral and imaging findings in our model correlated strongly with each other and have developed into a comprehensive set of outcome measures to be used in future therapeutic evaluations. Ongoing analyses are investigating the utility of PET ligands in this macaque model that bind to aggregated species of mHTT^66^. If successful, this will provide another imaging outcome measure to employ for preclinical therapeutic studies and which would be particularly beneficial in providing spatial resolution when assessing mHTT-lowering agents. Upon necropsy at 30 months post-surgery, we will also assess gray and white matter neuropathology using a variety of histological, molecular and biochemical techniques which will be correlated to behavioral and imaging findings at the same timepoint.

An advantage of viral-mediated HD macaques over the transgenic macaque models that have been created to date is that they can be generated in large enough numbers to appropriately power studies investigating novel biomarkers of disease progression and promising therapeutics. While the transgenic HD macaque model (exons 1-10 of the *hHTT* gene with 67-73Q) created by Chan and colleagues showed several key disease symptoms (progressive motor phenotypes, evidence of impulsivity and changes in temperament), along with white matter microstructural changes, striatal atrophy and aggregate formation, they only generated and characterized a small cohort of three transgenic animals, along with controls, precluding evaluation of any therapeutics.^67–71^ In addition to being able to generate larger cohorts of animals, viral-vector based macaque models allow for freedom in model design including the ability to assess different promoters, HTT fragment lengths, CAG repeat lengths, AAV serotypes (and combinations thereof) in a shorter timeframe, and with less expense, compared to transgenic and knock-in NHP models. For example, if desired, a longer prodromal phase in this AAV-based macaque model may be achievable by expanding the HTT fragment length (ie expressing the first 10 exons of human *mHTT* cDNA versus the first three) or shortening the CAG repeat number, particularly given that human HD patients show a direct correlation between CAG repeat length and age at disease onset. ^72^ Increasing the *mHTT* fragment length in this model would also expand the real estate available for *mHTT*- lowering therapeutics. Another possibility is to engineer patient-relevant single nucleotide polymorphisms into the mHTT transgene for evaluation of allele-specific HTT-lowering therapeutics.

As some of the HTT-lowering therapeutics currently under preclinical investigation are delivered using viral vectors (i.e. miRNAs, zinc finger protein-repressors, CRISPR-mediated gene editing, etc.) it will be critical to investigate the potential immune considerations of AAV-based therapeutics in this model, where neutralizing antibodies generated from creating the model could potentially reduce the efficacy of a second AAV vector to deliver its gene cargo. Should this be an issue, plasmapheresis, immunosuppression and/or capsid serotype switching using AAVs from distinct clades may be viable options. Other therapeutics also amendable to testing in this model include antisense oligonucleotides, stem cell-based therapeutics, small molecules, neurotransmitter-modulating pharmacotherapeutics, among others. We are hopeful that this new macaque model will become a widely used resource for the HD research community, both for identifying novel biomarkers of disease progression as well as testing promising therapeutic candidates.

## METHODS

### EXPERIMENTAL MODEL AND SUBJECT DETAILS

#### Animals

This study included 17 adult *Rhesus Macaques (Macaca mulatta)* (age 6-13; n=12 female, n=5 male) (**Table 1**). Monkeys were pair housed on a 12-hour light/dark cycle, provided with monkey chow rations twice daily, and given ad libitum access to water. Animal weights were recorded monthly by veterinary staff and heath checks were conducted by Oregon National Primate Research Center (ONPRC) veterinary and technical staff daily. The Institutional Animal Care and Use Committee and the Institutional Biosafety Committee at the ONPRC and Oregon Health and Science University (OHSU) approved all experimental procedures, and all of the guidelines specified in the National Institutes of Health Guide for the Care and Use of Laboratory Animals ^73^ were strictly followed.

#### Viral Vector Production

pAAV2.retro capsid plasmids were provided by the Karpova lab at the Howard Hughes Medical Institute (HHMI), Janelia Research Campus and pAAV2 capsid plasmids were supplied by the OHSU/ONPRC Molecular Virology Support Core. Plasmids containing the N171 N-terminal fragment sequence of human *HTT* bearing either 85 CAG repeats (HTT85Q, pathological *HTT* fragment) or 10 CAG repeats (HTT10Q, control *HTT* fragment) were manufactured by GenScript and subsequently cloned into a transgene cassette flanked by viral inverted terminal repeats (ITRs). *HTT* transgene expression was driven by a CAG promoter (cytomegalovirus (CMV) enhancer fused to the chicken beta-actin promoter) in both vector constructs. Recombinant AAV2 and AAV2.retro vectors were produced by the OHSU Molecular Virology Support Core and were prepared using a scalable transfection method as previously described by Weiss, et al. ^28^. Viral titers were determined by quantitative PCR of purified vector particles using a CAG primer/probe set: Forward: 5′-CCATCGCTGCACAAAATAATTAAAA-3′, Reverse: 5′-CCACGTTCTGCTTCACTCTC-3′, Probe: 5′-CCCCTCCCCACCCCCAATTTT-3′. Neutralizing Antibody (Nab) assays were carried out as previously reported ^28^, and animals with anti-AAV2 Nab titers of 1:20 or less were selected as study participants.

#### Surgery

Pre-surgical procedures and surgical parameters were identical to those previously reported ^28^, with a few noted deviations. Briefly, animals received 4 bilateral injections (8 injections total) of either a 1:1 mixture of AAV2 and AAV2.retro at a titer of 1e12 vg/mL (2e12 vg/ml combined) or a buffered saline injection w/ F-Pluronic (n=6 animals injected with AAV2-HTT85Q + AAV2.retro-HTT85Q, n=6 with AAV2-HTT10Q + AAV2.retro-HTT10Q and n=5 with phosphate buffered saline). Injections were made bilaterally into the pre-commissural caudate (90 ul) and putamen (95 ul) and into the post-commissural caudate (60 ul) and putamen (85 ul), for a volume of 330 ul per hemisphere. Following completion of all injections, dura was sutured closed, craniotomy sites were filled with gel foam, musculature and skin were sutured, and post-operative care was administered as previously described ^28^.

#### 3-Choice Spatial Delayed Response (SDR) Task

The 3-Choice Spatial Delayed Response assesses spatial working memory and was conducted in a Wisconsin General Testing Apparatus (WGTA) equipped with a plexiglass 3-well stimulus tray (6cm in diameter, 2.5cm depth, 8.5cm apart) (**Figure 1A**). Wells were arranged in a single row on the tray and were covered with identical plastic discs. At the beginning of each trial, an experimenter who was blind to the monkey’s treatment group uncovered one of the wells, placed a preferred food reward in the well, and replaced the cover on the well. The screen was then lowered and a stopwatch started. After a delay, the screen was raised, and the monkey was allowed to displace one of the well covers. If the baited well was chosen, the monkey was allowed to retrieve the food reward, after which the experimenter lowered the screen and marked the trial correct. If the monkey uncovered one of the non-baited wells, the screen was quickly lowered, the trial marked incorrect, and a new trial started. Prior to surgery, the animals were trained to complete this task to an 80% correct criterion using a 1-sec delay. 15 of 17 animals successfully acquired the rules of the task (**Table 1**). At the baseline, 3-, 6-, 9-, 14-, and 20-month study timepoints, monkeys completed two sessions of this task, on two consecutive days, with variable delays of 1s, 3s and 5s (8 trials each), resulting in 48 trials total. Each response was recorded and an inter-trial interval (ITI) of 15sec was initiated before advancing to the next trial.

#### Delayed Non-Match to Sample (DNMS) Task

The Delayed Non-Match to Sample (DNMS) task measures object recognition and familiarity. It was conducted in a soundproof testing chamber equipped with an automated testing apparatus. This apparatus included a 3M Microtouch Touch Screen monitor, a MedAssociates mini-M&M dispenser, and a speaker system. To begin this task, subjects were presented with a sample image in the middle of the screen. After touching the stimuli, the screen was cleared and a delay was initiated. After the delay, the sample image reappeared on either the left or right side of the screen along with a novel image (drawn from a library of over 5000 clipart images). Selection of the novel, non-matching, image resulted in a food reward, a positive tone and a 1s inter-trial interval (ITI), while selection of the familiar image resulted in a negative tone, no reward and a 5s ITI. Prior to surgery, animals were trained to complete this task to an 80% correct criterion using a 1-sec delay. 11 of 17 animals successfully acquired the rules of the task (**Table 1)**. At the baseline, 3-, 6-, 9-, 14-, and 20-month study timepoints, subjects completed two sessions, on two consecutive days, of 90 trials per session with intermixed delays (n=30 1s, n=30 5s, n=30 10s), resulting in 180 trials total.

#### Lifesaver Retrieval Task

Fine motor skill capability was assessed using the Lifesaver Retrieval task. This task was conducted in a WGTA equipped with a plexiglass cage front containing two arm openings as well as a plexiglass stimulus tray with 2 vertical, metal posts (7.5cm high, 11.5cm apart) on the left and right side of the tray **(Figure 1D)**. At the beginning of each trial, the experimenter threaded a Lifesaver candy onto one of the two posts. The screen was lifted to allow the monkey to retrieve the candy with their ipsilateral hand (right hand/right post, left hand/left post). For each session, the position of the Lifesaver alternated between the left and right post for 10 trails (n=5 left, n=5 right). Prior to surgery, all 17 animals successfully learned to complete this task (**Table 1**). Subjects completed two sessions of this task on two consecutive days at each timepoint (baseline, 3-, 6-, 9-, 14- and 20- month). The retrieval latency (recorded in seconds) was measured frame-by-frame using Adobe Premiere software. The retrieval latency was defined as the time interval between when monkey’s hand first passed through the fiberglass holes in the stimulus tray and ended when their hand was completely withdrawn back into the cage.

#### Neurological Ratings

An NHP-specific neurological rating scale (NRS) was used to score motor behaviors. Using methods previously described^74^, trained members of the lab who were blind to the experimental treatment groups observed each monkey for 30-45 minutes in their home environment. Each behavioral phenotype on the NRS was scored between 0-3, with a score of 0 representing normal behavior and a score of 3 representing severely abnormal behavior (see **Table S3** for a list of each behavior scored). Monkeys were rated at baseline, prior to surgery, and at each month post-surgery, with the addition of a 2-week post-surgery timepoint. Ratings were conducted during the same time of day, between 12pm-4pm. Additionally, we rated the animals using the NRS both prior to, and for 45 minutes following administration of the dopamine agonist, Apomorphine HCl (0.3 mg/kg, Sigma). Apomorphine ratings were conducted at baseline and 3-, 6-, 9-,14-, and 20-month post-surgery.

#### MRI Acquisition

All monkeys were anesthetized for the duration of the scanning session to prevent motion artifacts and to ensure their safety. Anesthesia was induced with Ketamine HCl (15mg/kg IM), and maintained via inhalation of 1-2% isoflurane gas vaporized in 100% oxygen. Animals were positioned on the scanner bed in a head-first supine orientation and immobilized in the head coil with foam padding. A fiducial marker (vitamin E tablet), was taped to the right side of the head prior to each scan. Blood oxygenation and heart rate were continually monitored throughout the scan by trained veterinary staff. Post scan, the animals were extubated, returned to their housing environment, and their recovery monitored closely for several hours by laboratory and veterinary staff.

A Siemens Prisma whole body 3T MRI system (Erlangen, Germany) with a 16-channel pediatric head rf coil was used to acquire all of the MR images. Four types of images were acquired: 3D T1-weighted magnetization-prepared rapid gradient-echo (MP-RAGE) ^75^, 3D T2-weighted sampling perfection with application optimized contrasts using different flip angle evolution (SPACE) ^76^, diffusion tensor imaging (DTI) scans, and resting state functional connectivity (rs-fMRI) scans. Details of the acquisition parameters for the structural (T1w/T2w) and DTI have been previously described and were identical to those reported by Weiss, et al. ^40^. Briefly, 3D SPACE imaging sequences were acquired with 0.5mm isotropic voxels (TE/TR = 385/3200 ms, flip angle = 120°, 320 x 320 x 224). Three SPACE images were acquired in each session (total acquisition time 29 minutes 42 seconds). For 3D MP-RAGE imaging sequences, voxel sizes and the field of view were identical to the 3D SPACE images (TE/TR/TI = 3.44/2600/913 ms, flip angle = 8°). Similarly, three MP-RAGE images were acquired in each imaging session (total acquisition time 31 minutes 9 seconds). Diffusion-weighed volumes were acquired using a spin-echo planar imaging (EPI) sequence with 1.0 mm isotropic voxels and TR/TE = 6700 ms/73 ms, GRAPPA factor = 2, echo train length = 52. Seven repetitions of 6 b0 volumes and 30 DW volumes with single b = 1000 s/mm^2^ and an anterior-to-posterior phase-encoding direction were acquired each session. To correct for susceptibility-induced distortions, a single b0 volume with a reversed (posterior-to-anterior) phase-encoding direction was also acquired (acquisition time 29 minutes 39 seconds). For rs-fMRI, BOLD images were acquired using T2-weighted GE-EPI sequence (TE/TR = 25/2290 ms and flip angle = 79°). Voxel sizes were 1.5 isotropic, and 784 3D volume images were collected. To control for effects of anesthesia on the BOLD signal, each rs-fMRI run started 45-50 min after the subject was first anesthetized with ketamine, and all animals were maintained on a constant level of 1% isoflurane during rs-fMRI acquisition. Following the rs-fMRI scan, there was a short reverse phase-encoded rs-fMRI scan (with 20 volume images) that was acquired for distortion correction (acquisition time 31 min 40 seconds).

#### SPACE/MP-RAGE Processing

SPACE and MP-RAGE images were processed with identical procedures. First, each of the three images collected during the imaging session were averaged. To accomplish this, the first scanned image from each modality was selected as the reference, and the other two images were registered to the reference with rigid-body transformations using ANTS (version 2.1, http://stnava.github.io/ANTs/) These warped images were subsequently averaged using FSL to produce merged T1w and T2w whole head images (version 5.0, http://fsl.fmrib.ox.ac.uk/fsl/fslwiki/). Due to similarities in the image contrasts of the T2w, b0, and EPI sequences, registration between the imaging modalities is superior when T2w data are used for structural alignment, compared to T1w data ^40, 77^. Additionally, data from human clinical populations has also demonstrated that T2w images provide improved sensitivity for VMB/TBM ^78^. Therefore, we used the ONPRC18 T2w template as our primary anatomical reference. For brain extraction, the merged T2w image was registered to the ONPRC18 T2w whole-head template ^40^ with b-spline non-linear transformations implemented in ANTS/FSL. Using the resulting registration parameters, the template brain mask was inversely mapped to the T2w merged images, verified by trained observers, and the brain extracted using FSL. Next, intensity bias correction was performed using the “N4BiasFieldCorrection” tool in ANTS ^79^. Finally, to co-align the structural scans, the merged T1w image was registered to the merged T2w image with a rigid-body transformation and skull stripped using the same mask. The merged T2w images were next aligned to ONPRC18 template space using b-spline non-linear transformations in ANTs/FSL, and resulting the Inverse Warp fields (representing the deformation from individual space to ONPRC18 template space) were used to calculate log Jacobian Determinant maps with the ANTs software package. Here, positive values denote areas where individual space was dilated to align with ONPRC18 space (and was therefore smaller), and negative values denote areas where individual space was contracted to align with ONPRC18 space (and was therefore larger). The log Jacobian Determinant maps from each subject at each timepoint were then analyzed using voxel wise-statistical comparisons implemented in FSL-SwE, described below.

#### DTI Processing

First, a susceptibility-induced off-resonance field (h) was calculated from the 6 pairs of b0 volumes with opposite phase-encoding direction using “topup,” included in the FSL library (https://fsl.fmrib.ox.ac.uk/) ^80^. A brain mask was manually generated for the resulting unwarped b0 image, and applied to the diffusion weighted (DW) volumes in a de-noising step that was implemented in MATLAB (script provided by Dr. Sune Jespersen, Aarhus University)^81^. To account for motion, the eddy current induced off-resonance field (e) and rigid-body transformations (r) between DW volumes were estimated simultaneously using “eddy” (FSL). Then, the 3 transformations (h, e, and r) were combined into one warp field to correct the de-noised DW volumes ^82^. Finally, the DTI-TK toolkit was used to fit the de-noised, eddy corrected b0s and DW volume to a single tensor (DTI) model. The DTI tensor maps for each animal at each timepoint were next aligned to the ONPRC18 tensor template with b-spline non-linear registrations, and parameter maps for FA, AD, RD, and MD were subsequently generated in template space using DTI-TK and then compared using voxel wise-statistical comparisons implemented in FSL-SwE, described below.

#### Resting-State fMRI

Preprocessing rsfMRI data was implemented with scripts from the Analysis of Functional Neuro Images (AFNI) Software package (https://afni.nimh.nih.gov) that were modified in-house for use with NHP data ^83^. The T2w SPACE brain images were used to co-register the resting-state data in anatomical space and transformed to ONPRC18 template space using the deformation fields generated by ANTs in the TBM pipeline described above. After regressing for signals of CSF and white matter, motion, and outliers’ censors ^84^, FSL-MELODIC software (https://fsl.fmrib.ox.ac.uk/fsl/fslwiki/MELODIC) was applied to the baseline data to derive group-level connectivity networks using Independent Component Analysis (ICA) ^85^. This approach identified 4 independent components (ICs) which appeared to align with previously described Resting-State Networks identified in macaques^86, 87^ and humans^85, 88–90^, overlapped primarily gray matter regions, and possessed low-frequency spectral power^91^ (**Figure S5)**. Based on these criteria, the remaining 4 ICs were not further examined^91^ **(Figure S6)**. To facilitate longitudinal group-level comparisons, dual-regression analysis was applied to each timepoint using FSL tools, focusing on the 4 ICs of interest. Z-scores from each of these 4 group-level ICs identified at baseline were then regressed against the individual 4D datasets at all of the timepoints to produce variance normalized time-courses for each of the 4 ICs for each induvial subject, and z-score maps indicating subject-level co-variance patterns with each of the 4 ICs brain-wide at each timepoint. These methods closely recapitulate the approaches used by Shnitko and Grant, (*personal communication*). Voxel-wise differences between groups were then analyzed using FSL-SwE, described below.

### QUANTIFICATION AND STATISTICAL ANALYSIS

#### Analysis of Behavioral Data

To account for individual differences in task performance pre-surgery, all behavioral measures (% correct, retrieval latency, NRS Score) were analyzed as change scores, calculated by subtracting the baseline values from each subsequent timepoint. Line charts and bar graphs were created using PRISM and IBM SPSS software packages illustrating group means (±SEM). Repeated measure ANOVAs, implemented in SPSS, were used to compare scores between groups over time. For the 3-Choice SDR task and the DNMS task, delay was included as a third factor in the ANOVA; for the Lifesaver Retrieval Task, hand was included as a third factor; and for the NRS, apomorphine was included as a third factor. Significance was defined by p<0.05, and post hoc tests with the Bonferroni correction for multiple comparisons were used to compare significant main effects. Additional planned comparisons were conducted using independent-sample t-test comparing between groups at each timepoint separately, to further elucidate the time course of any group differences.

#### Analysis of DTI, TBM, and rsfMRI data

For all imaging modalities (TBM, DTI, rsfMRI) parameter maps (log Jacobian Determinants, FA, AD, MD, RD, z-scores) were compared across timepoints using the FSL Sandwich Estimator (FSL-SwE) tool ^92^ with threshold free cluster enhancement (TFCE), 500 permutations ^93^, and a gray matter mask, a white matter mask, or a whole brain mask derived from the ONPRC18 labelmap. A threshold of p<0.01 was applied to all the results, however, due to the small sample sizes in this study, family-wise error correction was not applied. Visualizations of the thresholded p-value maps were created using FSLeyes. To summarize the regional distribution of significant voxel-wise changes, for each contrast we calculated the percent of p<0.01 voxels in each region of interest (ROI) in the ONPRC18 labelmap. This was accomplished by first using FSL tools to threshold and binarize and the p-value maps at p<0.01 and then AFNI tools (3dcalc) to extract volume information. Histograms were subsequently plotted using SPSS illustrating the percent volume of each ROI with significant changes from baseline. Finally, to assess the magnitude of the changes, ‘difference’ maps were calculated for each imaging parameter (log Jacobian Determinants, FA, AD, MD, RD, z-scores) by subtracting baseline from each subsequent timepoint (implemented in FSL). Then, for each parameter, masks were created that merged together the thresholded p-value maps from each timepoint, and the average value of each ‘difference’ map was calculated for the region under each p-value mask. This calculation was made brain-wide, rather than for each ROI separately. Using these data, changes were assessed with repeated measure ANOVAs including post-hoc group comparisons using the Bonferroni correction; additionally, these behavioral and imaging measures were collapsed across time (averaged) and correlated using 2-tailed Pearson correlations in SPSS.

### Data Availability

The ONPRC18 multimodal macaque brain atlas used in this manuscript has been deposited at the NeuroImaging Tools & Resources Collaboratory (https://www.nitrc.org/projects/onprc18_atlas) and is also published ^40^.

## ACKNOWLEDGEMENTS

We extend our sincere gratitude to the ONPRC Division of Animal Resources and Research Support for the excellent care provided to the rhesus macaques involved in this study, with special acknowledgement of the steadfast and expert veterinary efforts that Lauren Drew Martin, Theodore Hobbs, Melissa Berg, Brandy Dozier, Rob Zweig, Michael Reusz, Alona Kvitky, Kristy Ritchie, Amy Kujacznski and Isabel Bernstein contributed to this work.

## AUTHOR CONTRIBUTIONS

Conceptualization: J.L.M; Methodology: J.L.M, C.K., and A.R.W; Software: A.R.W, X.W., and Z.L; Formal Analysis: A.R.W, K.B., W.A.L, C.K., and J.L.M; Investigation: A.R.W, K.B., W.A.L., Z.L., X.W., and J.L.M; Writing – Original Draft: A.R.W.; Writing – Review & Editing: A.R.W, C.K, W.A.L., and J.L.M.; Funding Acquisition: J.L.M and C.K.

## FINANCIAL DISCLOSURE/COI

The authors declare no competing interests

## FUNDING SOURCES

NIH/NINDS R01NS099136, NIH/NINDS F32NS110149, NIH/NIA T32AG055378, NIH P51OD011092 and The Bev Hartig Huntington’s Disease Foundation.

## SUPPLEMENTARY FIGURES

**Figure S1.**
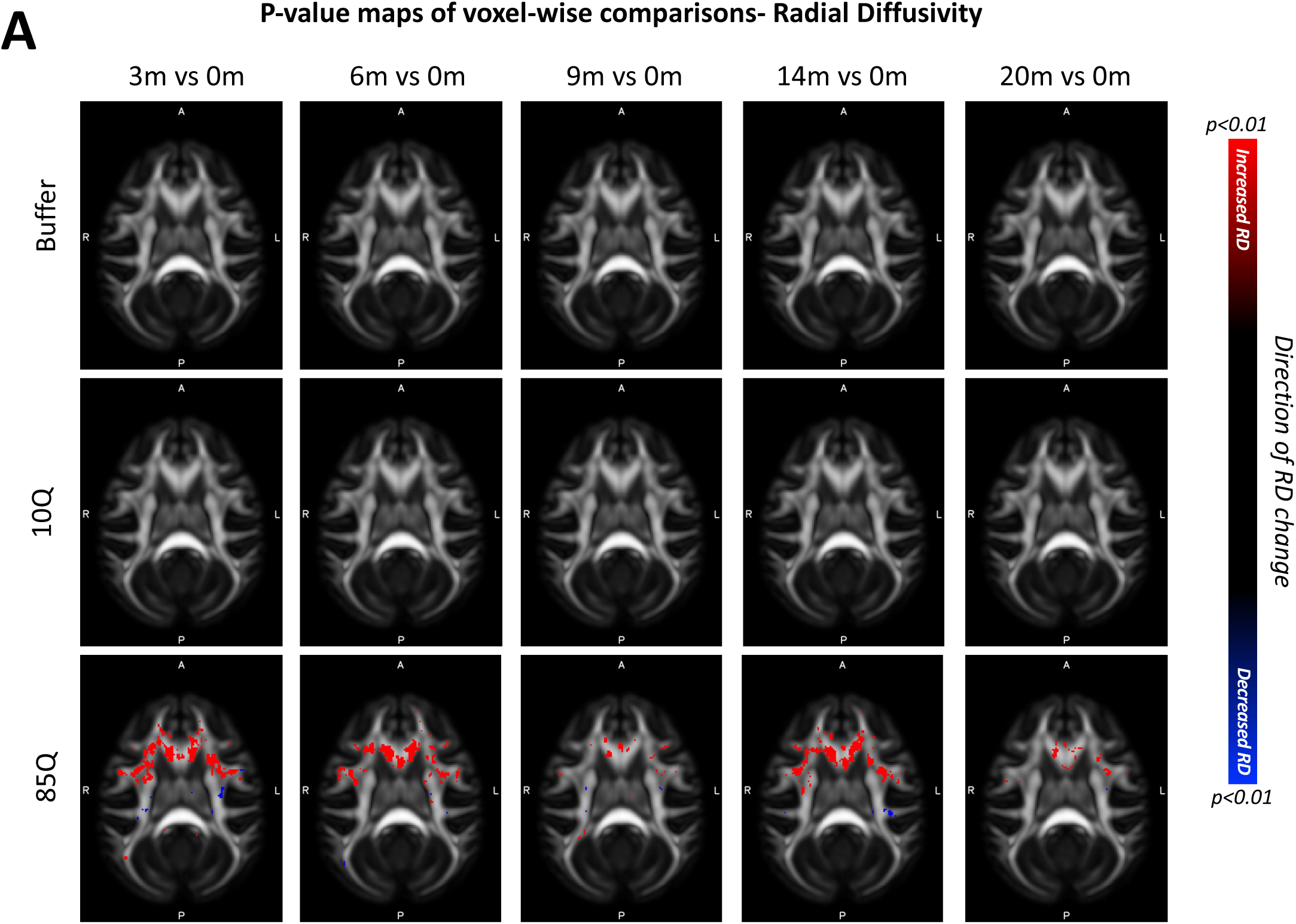
85Q-mediated changes in white matter Radial Diffusivity (RD). A) ONPRC18 FA template with overlaying p-value maps shown at a threshold of p<0.01. Blue voxels indicate areas of significant RD decrease, and red voxels indicate areas of significant RD increase. Although there were slight changes in the Buffer and 10Q over time, none of the contrasts reached statistical significance.

**Figure S2.**
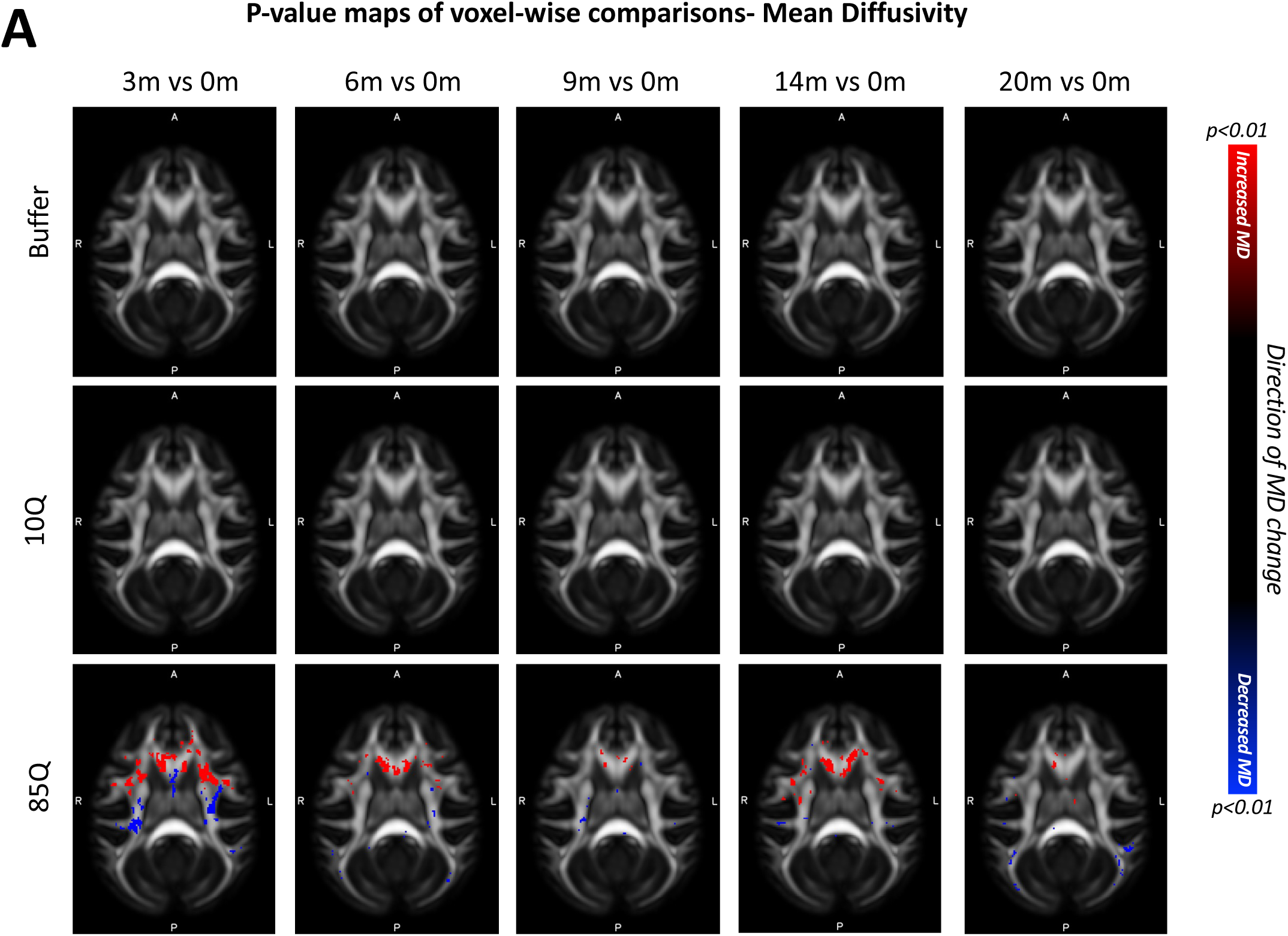
85Q-mediated changes in white matter Mean Diffusivity (MD). **A)** ONPRC18 FA template with overlaying p-value maps shown at a threshold of p<0.01. Blue voxels indicate areas of significant MD decrease, and red voxels indicate areas of significant MD increase. Although there were slight changes in the Buffer and 10Q over time, none of the contrasts reached statistical significance.

**Figure S3.**
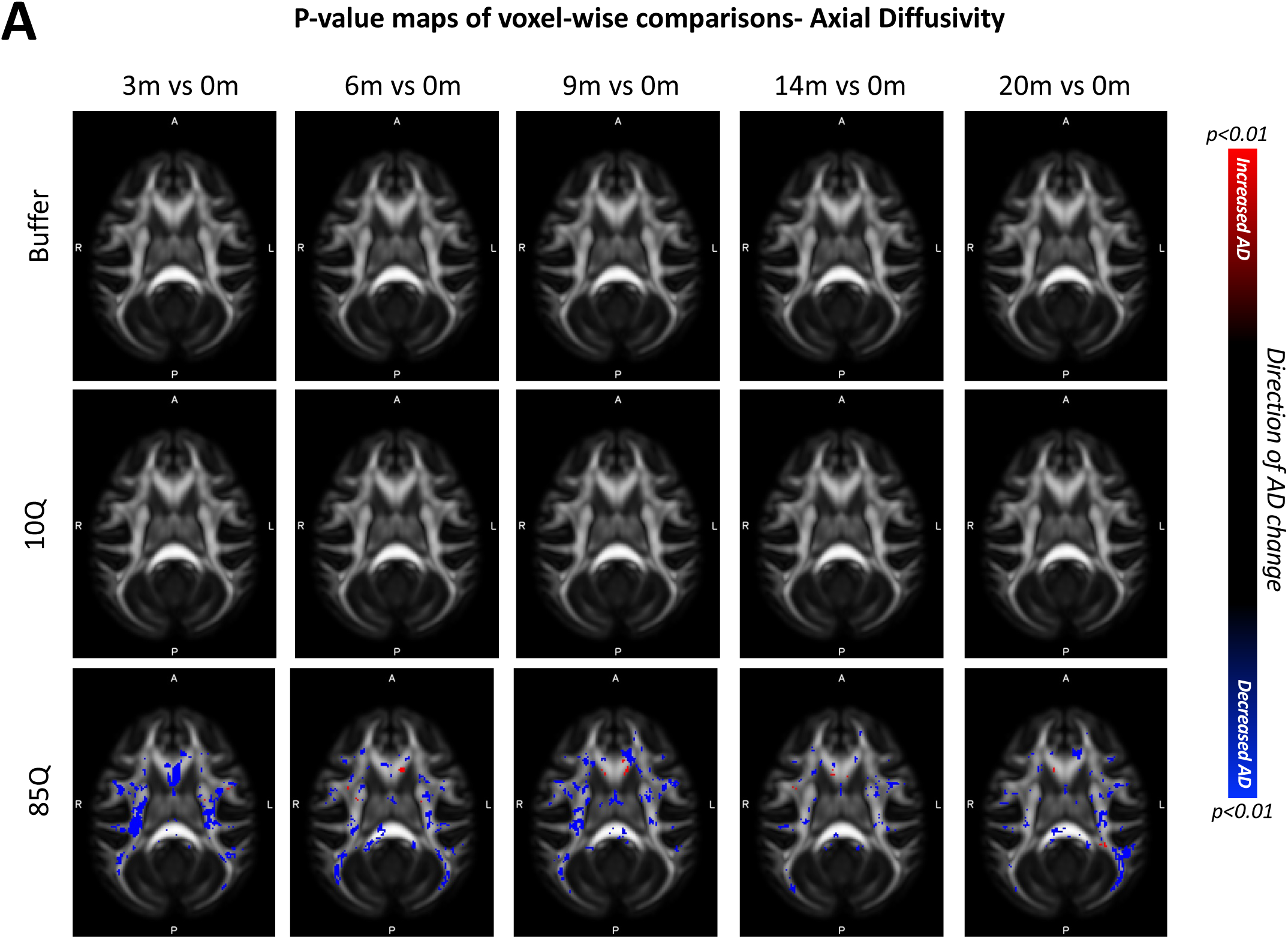
85Q-mediated changes in white matter Axial Diffusivity (AD). **A)** ONPRC18 FA template with overlaying p-value maps shown at a threshold of p<0.01. Blue voxels indicate areas of significant AD decrease, and red voxels indicate areas of significant AD increase. Although there were slight changes in the Buffer and 10Q over time, none of the contrasts reached statistical significance.

**Figure S4.**
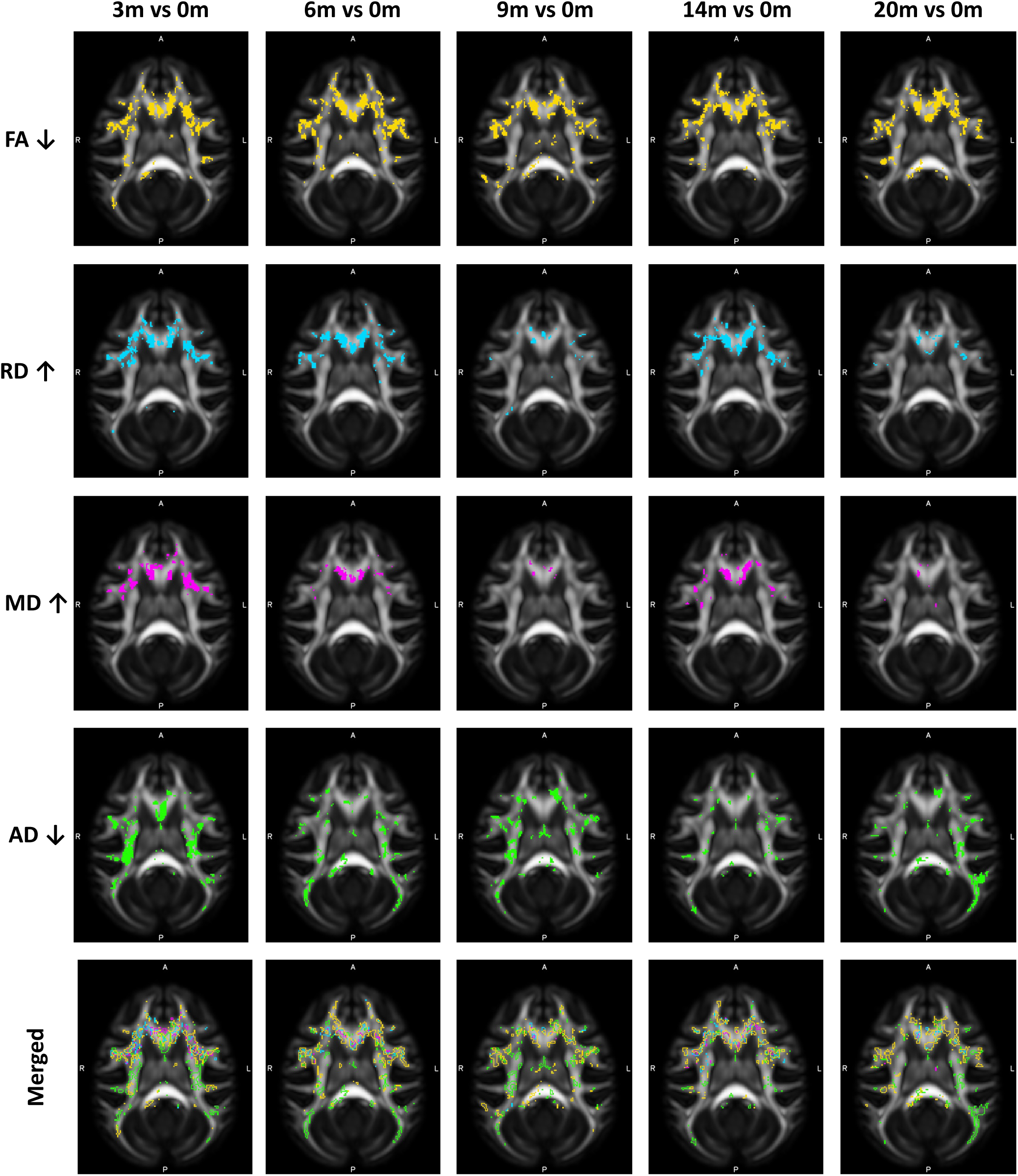
85Q-mediated changes in all diffusivity measures. **A)** ONPRC18 FA template with overlaying p-value maps shown at a threshold of p<0.01. Top row (yellow voxels) indicates areas of increased FA. Second row (turquoise voxels) indicate areas of increased RD. Third row (magenta voxels) indicate areas of increased MD. Forth row (green voxels) indicate areas of decreased AD. Bottom row shows the merged maps, to highlight areas of overlap. In general, FA decreases were closely aligned with RD increases and MD decreases.

**Figure S5.**
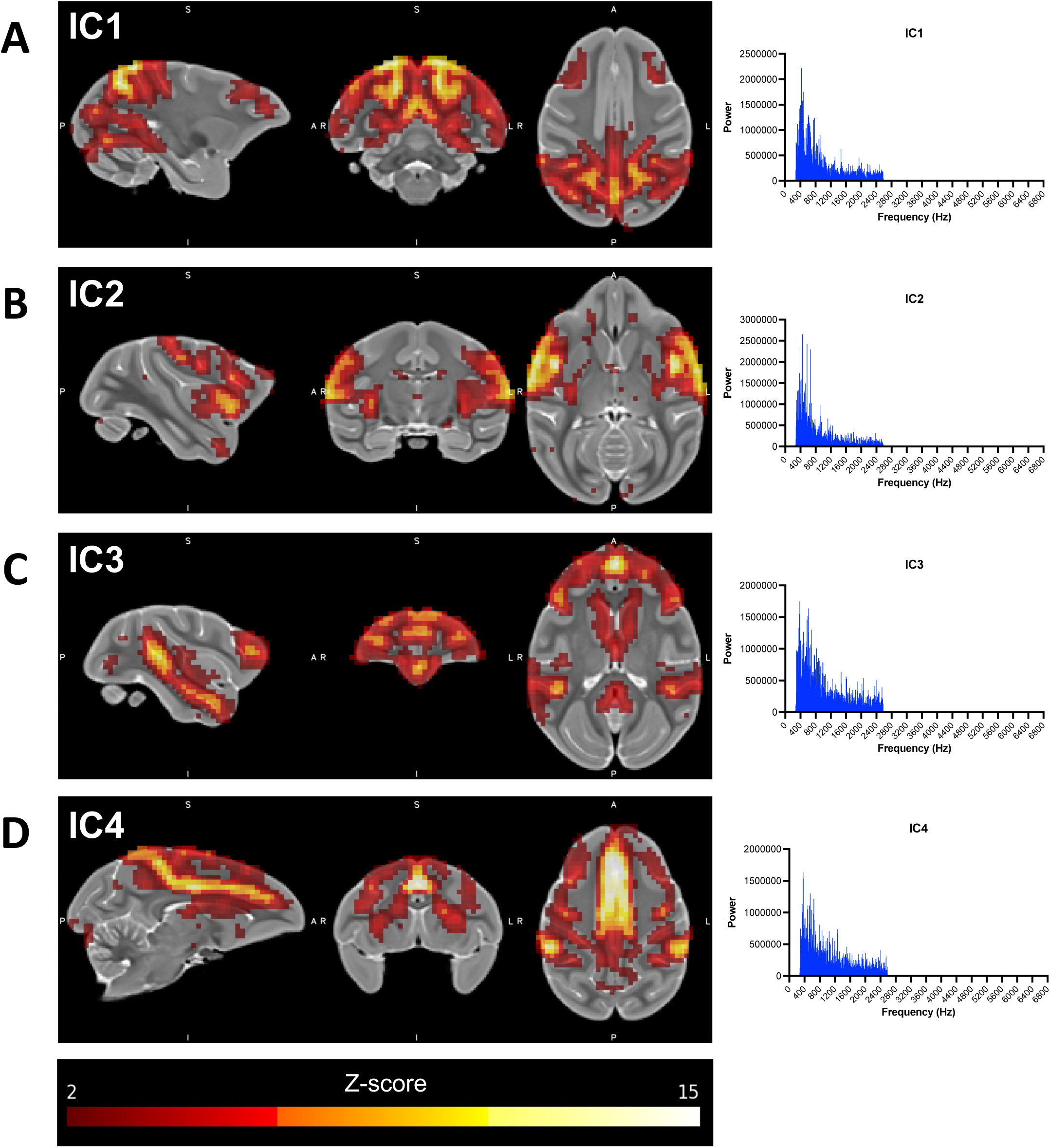
Resting-State Networks of Interest identified using Independent Component (IC) Analysis. (A-D) Illustrations showing the ONPRC18 T2w template with overlaying z-score maps from each IC of interest (shown at a threshold of z>2), along with histograms demonstrating the low-frequency spectral power associated with each^91^.

**Figure S6.**
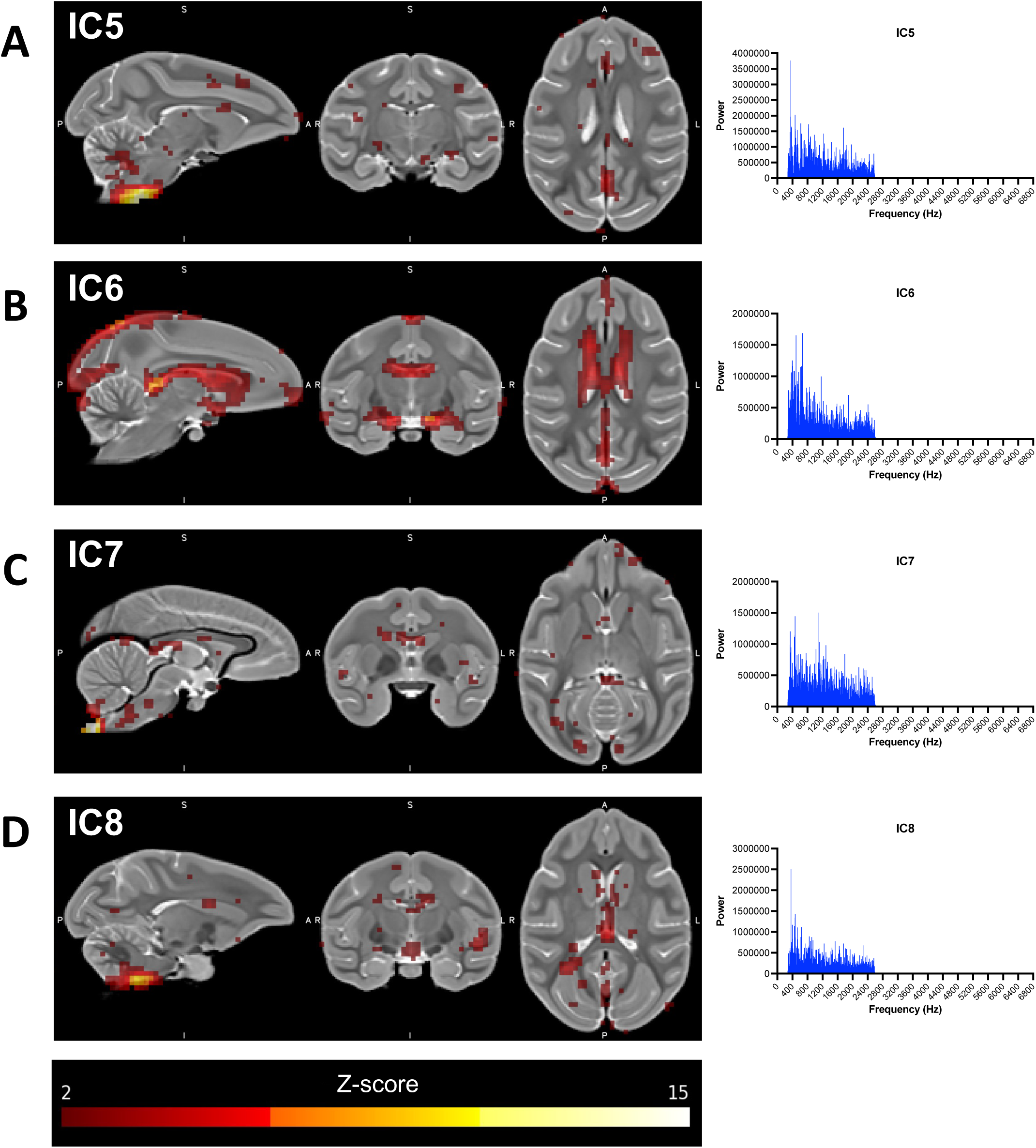
Networks identified using Independent Component Analysis associated with noise. (A-D) Illustrations showing the ONPRC18 T2w template with overlaying z-score maps from each omitted IC (shown at a threshold of z>2), along with histograms illustrating the relatively even distribution of spectral frequencies in each.

**Figure S7.**
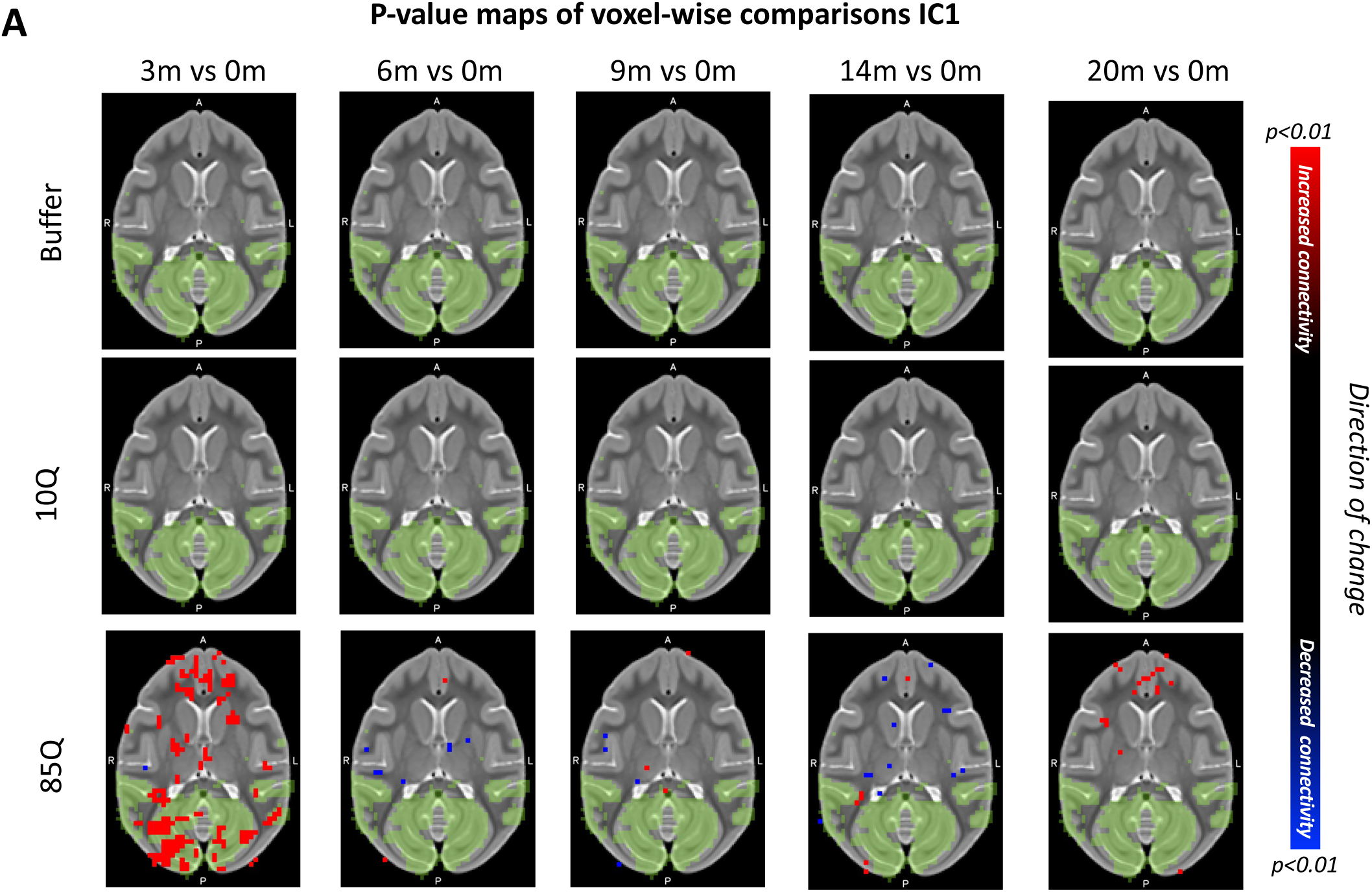
85Q-mediated changes Resting-State Functional Connectivity: IC1. **A)** ONPRC18 T2w template with overlaying p-value maps shown at a threshold of p<0.01. Blue voxels indicate regions of significant rsfc decrease, and red voxels indicate areas of significant rsfc increase. Green voxels indicate regions of IC1. Although there were slight changes in the Buffer and 10Q over time, none of the contrasts reached significance.

**Figure S8.**
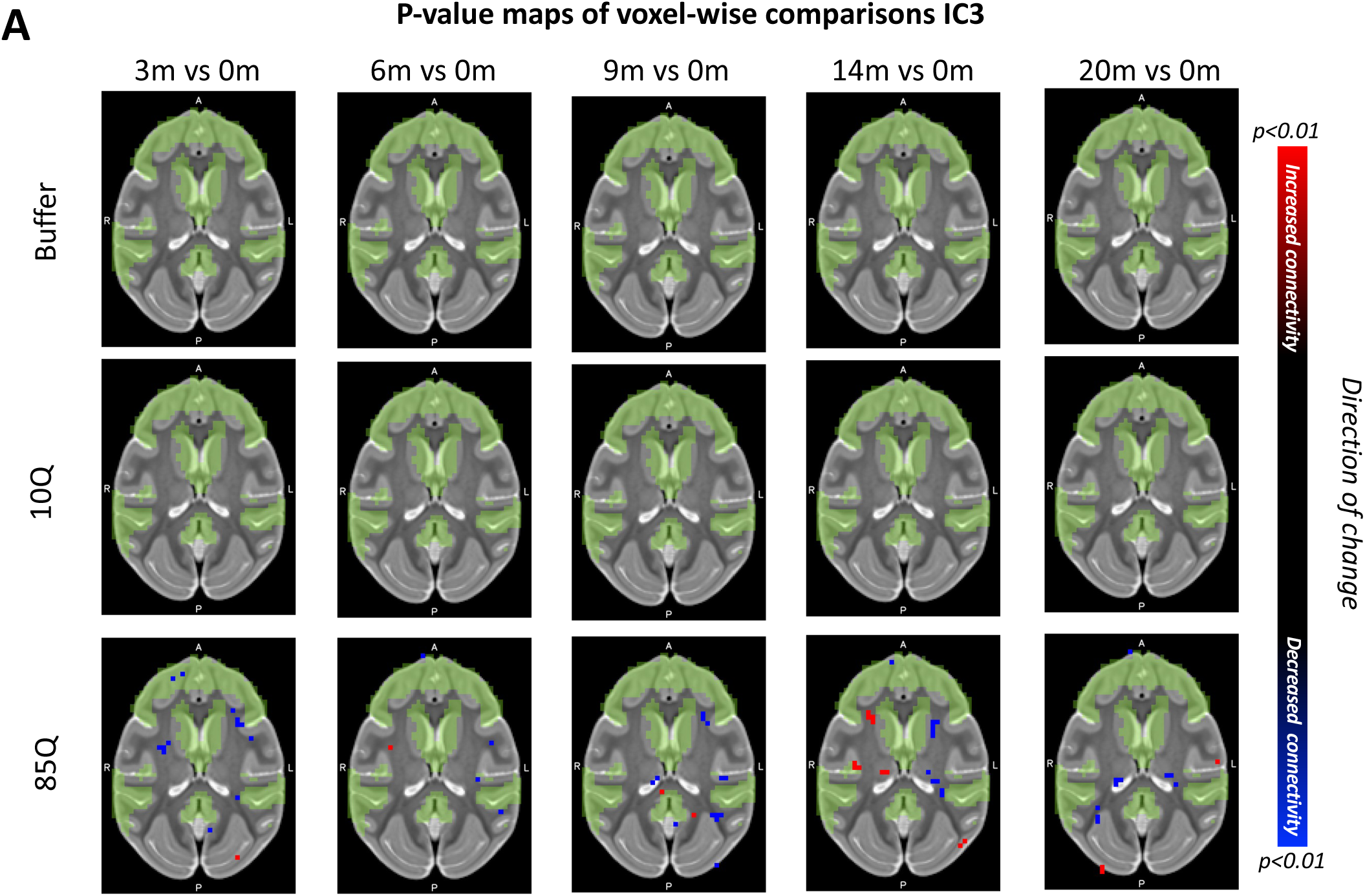
85Q-mediated changes Resting-State Functional Connectivity: IC3. **A)** ONPRC18 T2w template with overlaying p-value maps shown at a threshold of p<0.01. Blue voxels indicate regions of significant rsfc decrease, and red voxels indicate regions of significant rsfc increase. Green voxels indicate regions of IC3. Although there were slight changes in the Buffer and 10Q over time, none of the contrasts reached significance.

**Figure S9.**
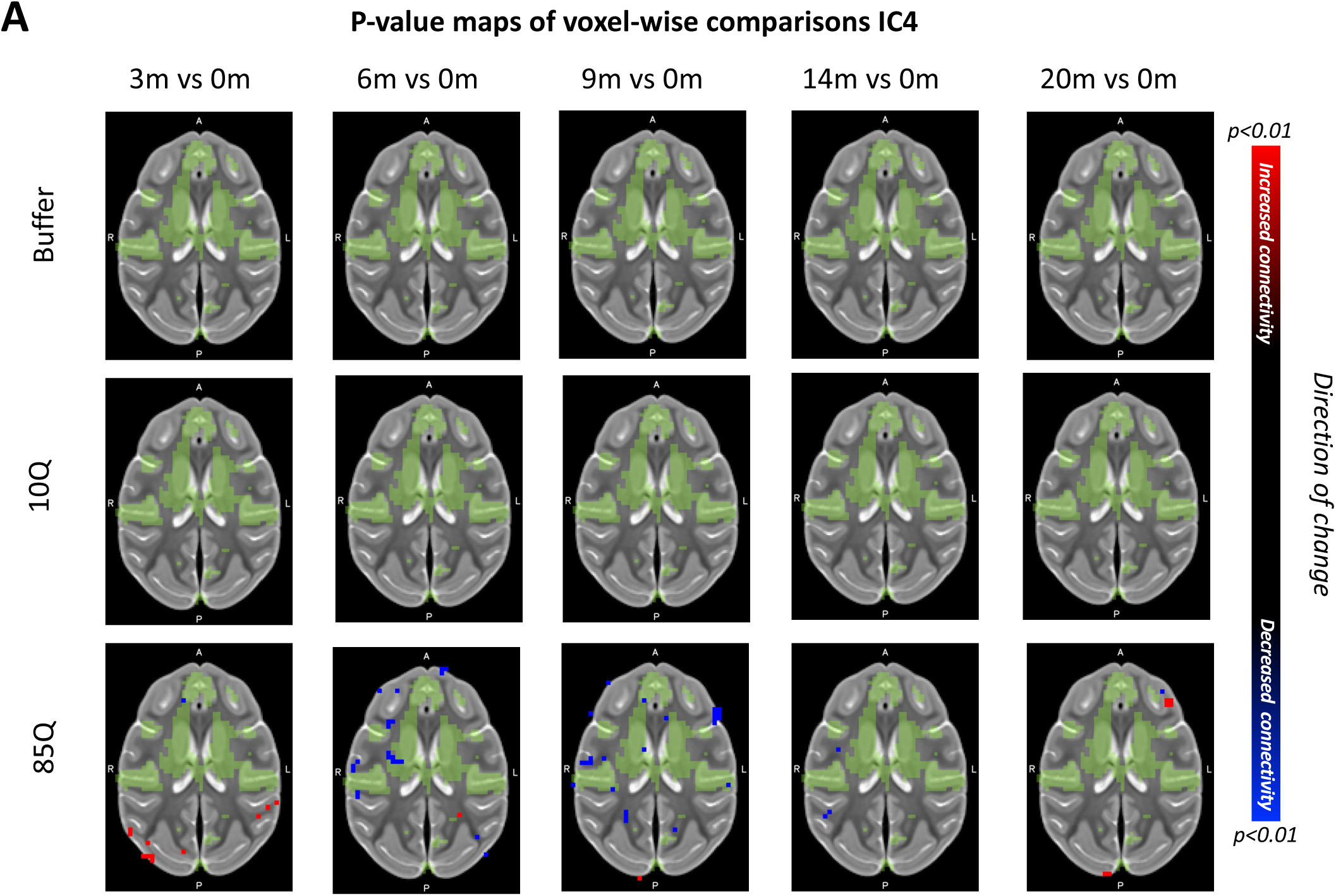
85Q-mediated changes Resting-State Functional Connectivity: IC4. **A)** ONPRC18 T2w template with overlaying p-value maps shown at a threshold of p<0.01. Blue voxels indicate regions of significant rsfc decrease, and red voxels indicate regions of significant rsfc increase. Green voxels indicate regions of IC4. Although there were slight changes in the Buffer and 10Q over time, none of the contrasts reached significance.

**Figure S10.**
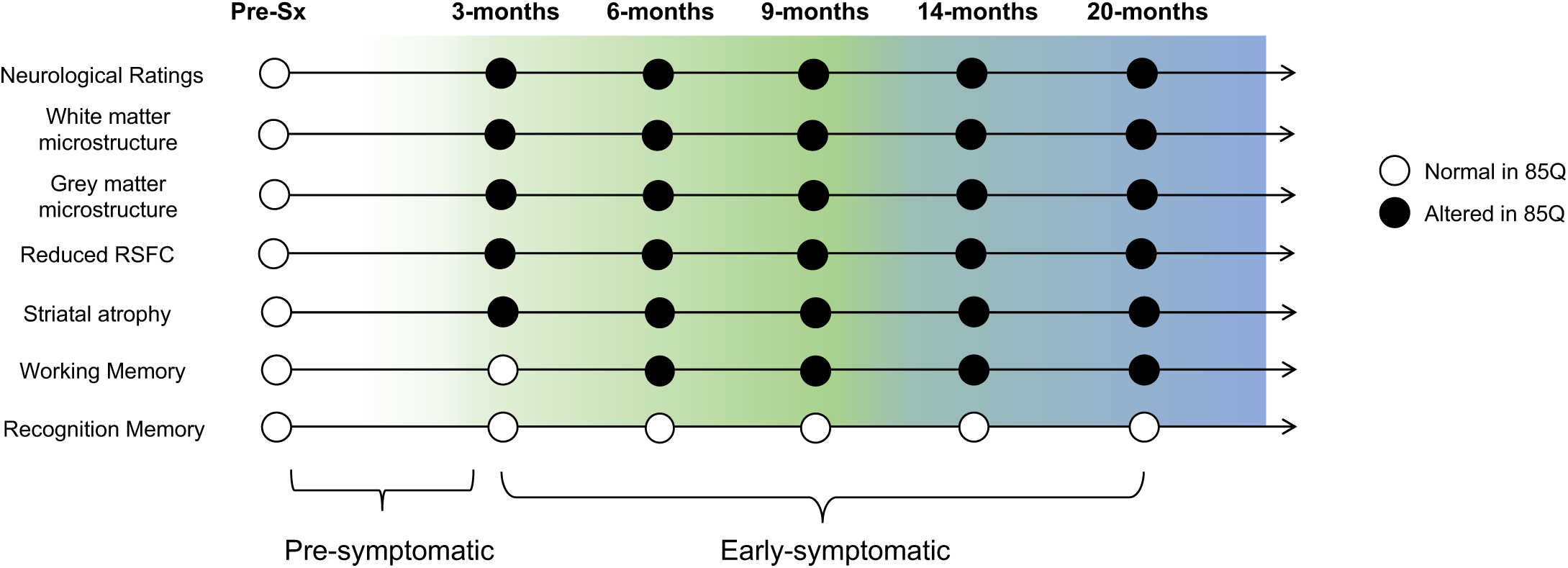
Timeline of findings in over the 20-month longitudinal study. Filled circles illustrate timepoints where group 85Q exhibited alterations from baseline measures, open circles illustrate timepoints where 85Q did not significantly differ from baseline.

## SUPPLEMENTARY TABLES

**Table S1.** Planned Group Comparisons for 3-Choice Spatial Delayed Response (SDR) task using Independent Sample T-tests. *p<0.05, **p<0.01, ***p<0.001

| Group Comparison | Timepoint | T-statistic | df | p-value |
| --- | --- | --- | --- | --- |
| 85Q vs 10Q | 3m | -1.189 | 9 | 0.265 |
|  | 6m | -1.981 | 9 | 0.079 |
|  | 9m | -3.196 | 9 | 0.011* |
|  | 14m | -4.886 | 9 | 0.001** |
|  | 20m | -2.194 | 9 | 0.056 |
| Buffer vs 10Q | 3m | -1.137 | 7 | 0.293 |
|  | 6m | -2.426 | 7 | 0.046* |
|  | 9m | -1.484 | 7 | 0.181 |
|  | 14m | -1.974 | 7 | 0.089 |
|  | 20m | -1.033 | 7 | .336 |
| Buffer vs 85Q | 3m | -2.047 | 8 | 0.075 |
|  | 6m | -3.652 | 8 | 0.006** |
|  | 9m | -5.762 | 8 | 0.0004*** |
|  | 14m | -6.043 | 8 | 0.0003*** |
|  | 20m | -3.392 | 9 | 0.009** |

**Table S2.**
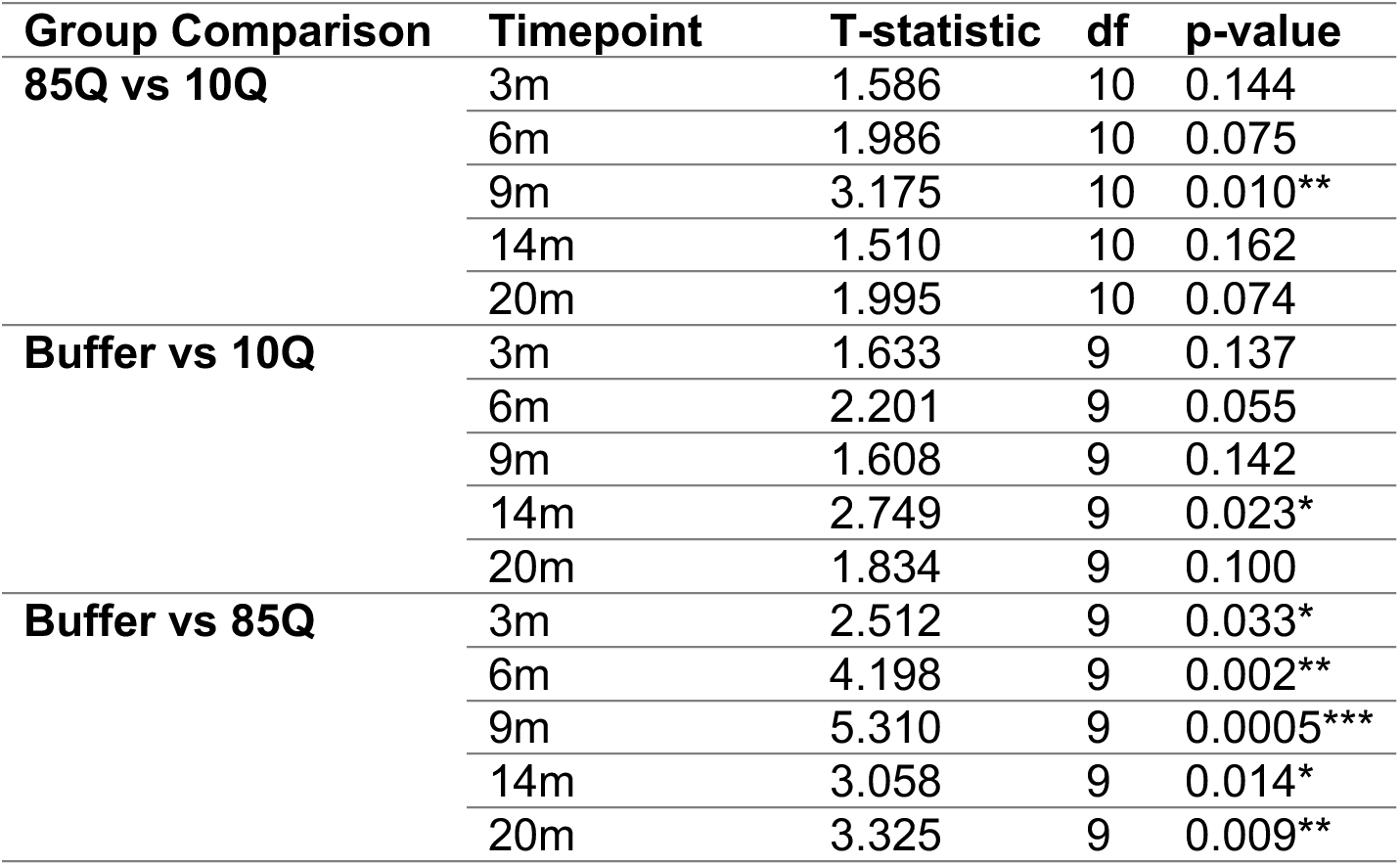
Planned Comparisons for Lifesaver Retrieval Latencies using Independent Sample T-tests. *p<0.05, **p<0.01, ***p<0.001

**Table S3.**
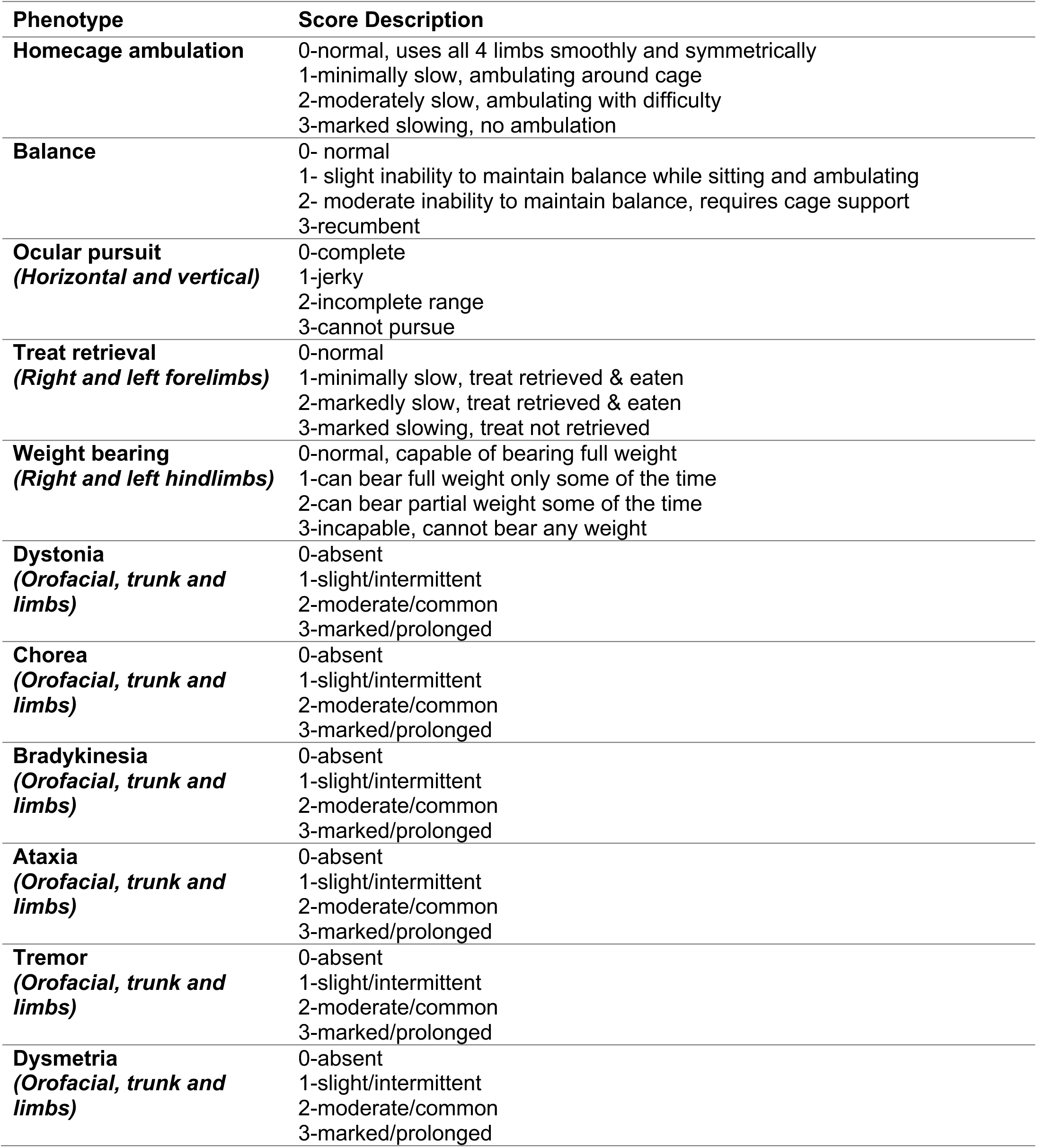
**Neurological Rating Scale for Nonhuman Primates**. Behaviors were scored cage-side by trained observers, blinded to treatment condition, during 30-45 minute focal observations. Scores were summed across categories to generate a total NRS score. Higher scores indicate more severe phenotypes.

**Table S4.** Planned Group Comparisons in Monthly NRS Scores using Independent Sample T-tests. *p<0.05, **p<0.01, ***p<0.001

| Group Comparison | Timepoint | T-statistic | df | p-value |
| --- | --- | --- | --- | --- |
| 85Q vs Buffer | 0.5m | -1.809 | 9 | 0.1039 |
|  | 1m | -1.52 | 9 | 0.1629 |
|  | 2m | 0.734 | 9 | 0.4819 |
|  | 3m | 2.555 | 9 | 0.0309* |
|  | 4m | 2.352 | 9 | 0.0432* |
|  | 5m | 3.371 | 9 | 0.0082** |
|  | 6m | 4.629 | 9 | 0.0012** |
|  | 7m | 2.961 | 9 | 0.0159* |
|  | 8m | 7.151 | 9 | 5.36E-05*** |
|  | 9m | 3.16 | 9 | 0.0115* |
|  | 10m | 4.931 | 9 | 0.0008*** |
|  | 11m | 5.149 | 9 | 0.0006*** |
|  | 12m | 8.613 | 9 | 1.22E-05*** |
|  | 13m | 5.056 | 9 | 0.0007*** |
|  | 14m | 6.945 | 9 | 6.72E-05*** |
|  | 15m | 8.906 | 9 | 9.30E-06*** |
|  | 16m | 5.002 | 9 | 0.0007*** |
|  | 17m | 5.859 | 9 | 0.0002*** |
|  | 18m | 12.043 | 9 | 7.47E-07*** |
|  | 19m | 11.687 | 9 | 9.64E-07*** |
|  | 20m | 6.182 | 9 | 0.0002*** |
|  | 21m | 11.442 | 9 | 1.15E-06*** |
| 85Q vs 10Q | 0.5m | 1 | 10 | 0.3409 |
|  | 1m | 1 | 10 | 0.3409 |
|  | 2m | 1.861 | 10 | 0.0924 |
|  | 3m | 2.996 | 10 | 0.0134* |
|  | 4m | 2.785 | 10 | 0.0193* |
|  | 5m | 3.478 | 10 | 0.0059** |
|  | 6m | 5.082 | 10 | 0.0005*** |
|  | 7m | 3.411 | 10 | 0.0066** |
|  | 8m | 6.742 | 10 | 5.09E-05*** |
|  | 9m | 3.034 | 10 | 0.0126* |
|  | 10m | 5.117 | 10 | 0.0005*** |
|  | 11m | 5.757 | 10 | 0.0002*** |
|  | 12m | 9.396 | 10 | 2.81E-06*** |
|  | 13m | 5.371 | 10 | 0.0003*** |
|  | 14m | 6.103 | 10 | 0.0001*** |
|  | 15m | 7.59 | 10 | 1.86E-05*** |
|  | 16m | 6.277 | 10 | 9.17E-05*** |
|  | 17m | 7.59 | 10 | 1.86E-05*** |
|  | 18m | 13.348 | 10 | 1.07E-07*** |
|  | 19m | 11.364 | 10 | 4.87E-07*** |
|  | 20m | 5.813 | 10 | 0.0002*** |
|  | 21m | 6.063 | 10 | 0.0001*** |
| Buffer vs 10Q | 0.5m | -2.018 | 9 | 0.074 |
|  | 1m | -1.764 | 9 | 0.112 |
|  | 2m | -1.543 | 9 | 0.157 |
|  | 3m | -0.905 | 9 | 0.389 |
|  | 4m | -0.905 | 9 | 0.389 |
|  | 5m | 0 | 9 | 1 |
|  | 6m | -0.905 | 9 | 0.389 |
|  | 7m | -0.905 | 9 | 0.389 |
|  | 8m | 0 | 9 | 1 |
|  | 9m | 0.267 | 9 | 0.796 |
|  | 10m | 0 | 9 | 1 |
|  | 11m | -0.408 | 9 | 0.693 |
|  | 12m | -0.905 | 9 | 0.389 |
|  | 13m | 0 | 9 | 1 |
|  | 14m | 0.905 | 9 | 0.389 |
|  | 15m | 0.267 | 9 | 0.796 |
|  | 16m | -1.398 | 9 | 0.196 |
|  | 17m | -0.811 | 9 | 0.438 |
|  | 18m | -0.129 | 9 | 0.9 |
|  | 19m | 0 | 9 | 1 |
|  | 20m | 0.267 | 9 | 0.796 |
|  | 21m | 1.945 | 9 | 0.084 |

**Table S5.** Planned Comparisons for Pre- vs Post-Apomorphine NRS Scores at each timepoint using Paired-Sample T-tests. *p<0.05, **p<0.01

| Group Comparison | Timepoint | T-statistic | df | p-value |
| --- | --- | --- | --- | --- |
| **Buffer** | 3m | -1.500 | 4 | 0.208 |
|  | 6m | -1.633 | 4 | 0.178 |
|  | 9m | -1.633 | 4 | 0.178 |
|  | 14m | -1.238 | 4 | 0.284 |
|  | 20m | -1.633 | 4 | 0.178 |
| **10Q** | 3m | 0.000 | 5 | 1.000 |
|  | 6m | -1.000 | 5 | 0.363 |
|  | 9m | -1.195 | 5 | 0.286 |
|  | 14m | -0.696 | 5 | 0.518 |
|  | 20m | -0.598 | 5 | 0.576 |
| **85Q** | 3m | -3.050 | 5 | 0.028* |
|  | 6m | -3.512 | 5 | 0.017* |
|  | 9m | -4.540 | 5 | 0.006** |
|  | 14m | -1.784 | 5 | 0.135 |
|  | 20m | -2.154 | 5 | 0.084 |

## Notes

https://www.nitrc.org/search/?type_of_search=group&q=ONPRC18

### Competing Interest Statement

The authors have declared no competing interest.

